# Phospho-proteomics identifies D-group MAP kinases as substrates of the Arabidopsis tyrosine phosphatase RLPH2

**DOI:** 10.1101/2024.08.26.609716

**Authors:** Anne-Marie Labandera, Ryan Toth, Sierra Mitchell, Jayde J Johnson, Juliette Puyaubert, Emmanuel Baudouin, R. Glen Uhrig, Greg B Moorhead

## Abstract

Despite being one of the few bona fide plant tyrosine phosphatases, RLPH2 has no known substrates. Utilizing phospho-proteomics, we identified the activation loop phospho-tyrosine of several D-group mitogen activated protein kinases (MPKs) as potential RLPH2 substrates. All Arabidopsis D-Group MPKs possess a TDY activation loop phosphorylation motif, whereas other MPKs (Groups A, B and C) contain a TEY motif. Our findings reveal that RLPH2 has a strong preference for aspartate (D) in the TXY motif, providing specificity for RLPH2 to exclusively target and dephosphorylate the D-Group MPKs. Additionally, D-Group MPKs contain a unique activation loop insertion that conforms to a protein phosphatase 1 (PP1) binding motif, with findings presented here confirming Arabidopsis PP1 phosphatases dock at this site. Intriguingly, only D-group MPKs among all identified Arabidopsis protein kinases possess this PP1 recruiting motif. Using multiple RLPH2 deficient plant lines, we demonstrate that RLPH2 represses seed dormancy release. Overall, this work highlights the power of phospho-proteomics in identifying substrates of this novel plant tyrosine phosphatase, while also revealing new complexities in the interactions between MPK activation loops and multiple phospho-mediated cell signaling events.

**One sentence summary:** Phospho-proteomic analysis reveals *Arabidopsis* tyrosine phosphatase RLPH2 dephosphorylates the activation loop of D-Group mitogen activated protein kinases.

## Introduction

Protein phosphorylation machinery, comprised of protein kinases and phosphatases, is highly conserved across eukaryotic organisms with protein phosphorylation recognized as a fundamental mechanism of controlling protein function. Proteins are primarily phosphorylated on the hydroxy amino acids serine (pS), threonine (pT) and tyrosine (pY) with eukaryotic phospho-proteomes being approximately 86%, 12%, and 2% on each of these residues, respectively (*1, 2*), including plants (*3*). The protein phosphatases that remove the phosphate from these protein sidechains are highly conserved across eukaryotes and are divided into four groups based primarily on primary amino acid sequence defining catalytic motifs and domains. These four groups include: the protein tyrosine phosphatases (PTP), the aspartate dependent phosphatases, the phosphoprotein phosphatases (PPP) comprised of PP1, PP2A, PP2B, PP4-PP7-like, and the Mg^2+^-or Mn^2+^-dependent protein phosphatase (PPM) family (*2, 4, 5*). Although the tyrosine phosphatases, based on their characterization in humans, display substrate specificity because of differing active site architecture, the serine/threonine specific enzymes from the PPP family are thought to achieve much of their specificity from associated regulatory subunits (*2, 4*). This is best characterized in the type one phosphatases (PP1), the majority of which dock the PP1 catalytic subunit via a short linear motif (SLiM) designated RVXF (*1, 2, 6*).

Like humans, plants maintain all four conserved protein phosphatase families, with the exception that plants have very few ‘mammalian-type’ protein tyrosine phosphatases, despite possessing tyrosine phosphorylation levels that parallel other eukaryotes (*3, 7, 8*). Recently we characterized the *Arabidopsis thaliana* (At) protein phosphatase RLPH2 (AtRLPH2), which based on its primary amino acid sequence is a PPP family serine/threonine protein phosphatase (like PP1), but possesses a clear preference for phosphotyrosine peptides as substrates when compared to phosphoserine/threonine peptides (*8*). AtRLPH2 selectively dephosphorylated the pY and not the pT of a human ERK1/2 activation loop peptide, and the presence of pT increased activity against the pY site (*7*). The mechanism underlying this specificity became clear when we solved the structure of AtRLPH2 (*7*). Uniquely, the active site of AtRLPH2 is much deeper compared to pS/pT specific phosphatases (e.g. PP1 or PP2A) allowing the larger pY side chain to reach the base of the pocket to be dephosphorylated, whereas pS/pT were modeled to be too short to allow for efficient dephosphorylation (*4*). Crystallization with the phosphate mimetic tungstate revealed that in addition to the active site, a basic pocket adjacent to the active site also bound tungstate, suggesting this was an additional site for binding a phospho-residue of substrates, much like the human dual specificity phosphatase VHR/DUSP3 (*7, 9*). Further modelling demonstrated that when a substrate pY peptide bound the AtRLPH2 active site, a pT two residues away was perfectly positioned to dock in the basic pocket. This configuration suggests that the basic pocket likely acts to recruit substrates to AtRLPH2 that are phosphorylated on threonine, facilitating dephosphorylation on a pY two amino acids C-terminal to pT, thus providing additional substrate specificity to the enzyme. Further analysis of plant phospho-proteome databases has uncovered multiple examples of pTxpY within protein sequences (table S1) and these could represent potential RLPH2 substrates.

To discover RLPH2 substrates, we undertook a quantitative phospho-proteomics approach. Knowing that biochemically RLPH2 is a tyrosine phosphatase, we isolated phospho-tyrosine containing peptides from the rosettes of Arabidopsis wildtype (WT) Nössen and two *rlph2* knockout lines. In the absence of AtRLPH2, proteomic analysis revealed a dramatic enrichment of tyrosine phosphorylated peptides derived from the activation loops of a subset of Arabidopsis mitogen activated protein kinases (MPKs), specifically D-Group MPKs. We then demonstrated that RLPH2 targets D-Group MPKs, discriminating between the TDY (D-Group MPKs only) versus TEY (all other MPKs) activation loop motifs. Additionally, we uniquely find that D-Group MPKs also dock protein phosphatase one (PP1) through a highly conserved PP1 binding RVXF SLiM. Limited biological roles have been attributed to D-Group MPKs, with D-Group MPK8 found to be involved in promoting seed germination in a gibberellic acid (GA) dependent manner (*10*). Correspondingly, our results indicate that AtRLPH2 is a negative regulator of germination in both an abscisic acid (ABA) and GA dependent manner.

## Results

### Phospho-proteomics to uncover AtRLPH2 substrates

To identify substrates of the tyrosine phosphatase AtRLPH2 we undertook a phospho-proteomic comparison between WT Nössen plants and two independent *rlph2* (*rlph2-1* and *rlph2-2*) mutant alleles. Biochemically, AtRLPH2 behaves as a tyrosine phosphatase (*7, 8*), making the goal of this comparison to elucidate the endogenous substrates of AtRLPH2 through the identification of tyrosine phosphorylation events present in *rlph2* versus WT plants. Here, we expect that the loss of AtRLPH2 should result in hyper-phosphorylation of endogenous substrates. To achieve this, we first enriched the global phospho-proteome of rosette tissue using TiO_2_, and then further enriched the tyrosine phosphorylated peptides from the global phospho-proteome by immunoprecipitation (IP) using anti-phospho-tyrosine (pY) specific antibodies (Fig. 1). Proteomic analysis of anti-pY IP eluates revealed tyrosine phosphorylated peptides belonging to the activation loop of three D group mitogen activated protein kinases (MPK9, 18 and 20) being specifically detected in the *atrlph2* lines, but absent in WT plants (Table 1). Notably, we found phosphotyrosine peptides corresponding to the activation loop of other MPK groups were enriched in both *rlph2* and WT plants suggesting a D-group specificity (table S2).

**Figure 1.**
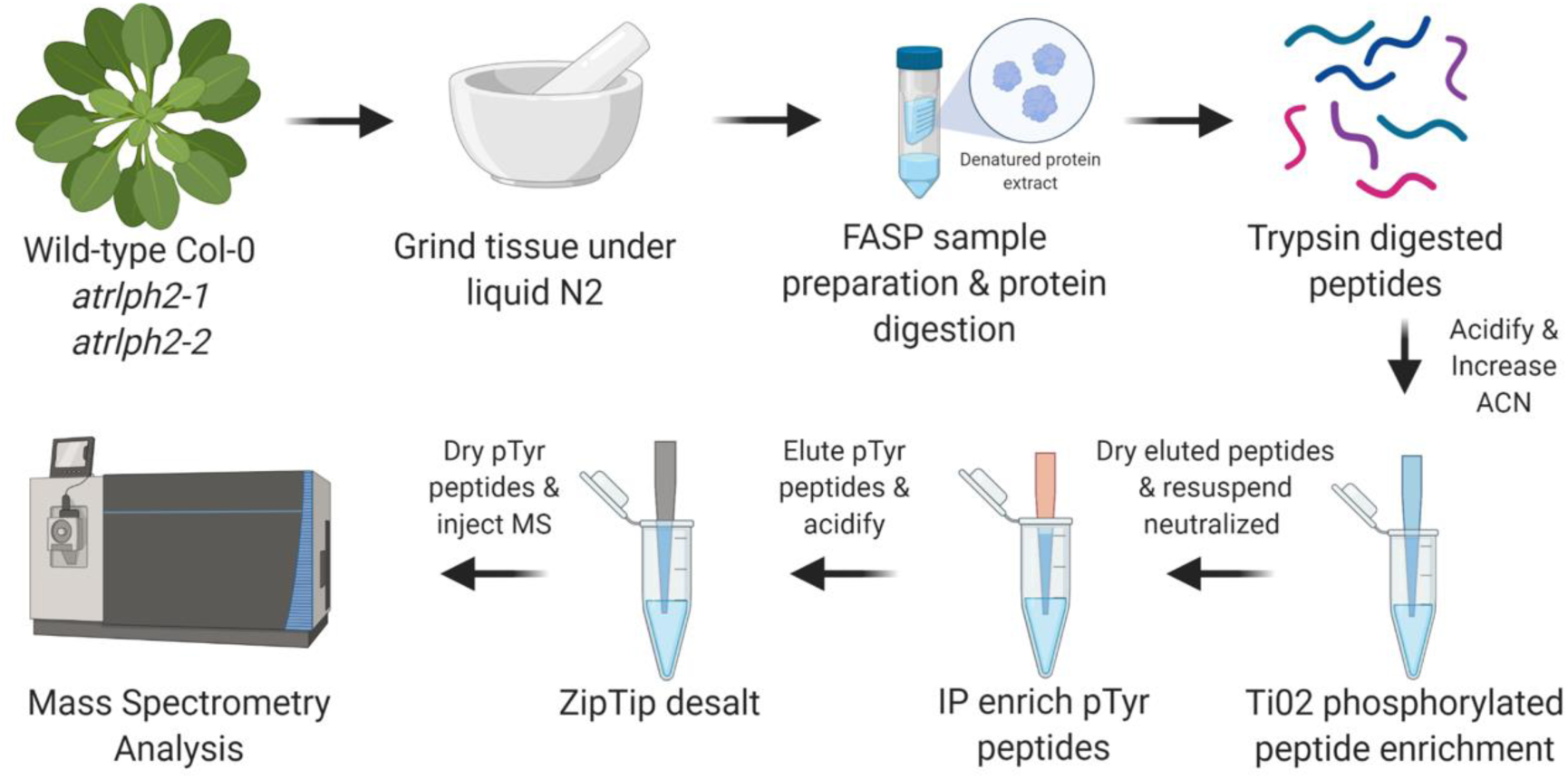
Schematic of the AtRLPH2 phospho-tyrosine (pY) substrate enrichment pipeline.

**Table 1.** AtRLPH2 preferentially dephosphorylates D-Group MAP kinases (MPKs). D-Group MAP kinases (MPKs) are exclusively phosphorylated in *atrlph2-1* and *atrlph2-2* relative to other MPKs. Shown are the tabulated results of two complete independent experiments in which phosphorylated-tyrosine (pY) containing peptides were isolated through a sequential Ti0_2_ enrichment / anti-pY immunoprecipitation workflow (Fig. 1). The complete list of all anti-pY immunoprecipitated phosphorylated proteins are found in table S2. Highlighted in red is the phosphorylated tyrosine residue (pY) identified, while identified oxidized methionine residues are denoted (oxM). The number of independent experiments in which each phosphorylated peptide is found (x/2) is shown under the corresponding genotype delimiter.

| AGI | Description | Phosphorylated Peptide | PTM<br>LOCALIZATION<br>PROBABILITY | MASCOT<br>SCORE | WT_Col-0 | *atrlph2-1* | *atrlph2-2* |
| --- | --- | --- | --- | --- | --- | --- | --- |
| AT2G42880 | ATMPK20, MPK20 \| MAP kinase 20 | VAFNDTPTTIFWTD(**pY**)VATR | 100% | 48.35 | 0/2 | 1/2 | 2/2 |
| AT3G18040 | ATMPK9 \| MAP kinase 9 | VSFNDAPSAIFWTD(**pY**)VATR | 100% | 39.53 | 0/2 | 1/2 | 2/2 |
| AT1G53510 | ATMPK18, MPK18 \| MAP kinase 18 | VAFNDTPTTVFWTD(**pY**)VATR | 69% | 35.27 | 0/2 | 0/2 | 2/2 |
| AT5G19010 | ATMPK16 \| MAP kinase 16 | VAFNDTPTAIFWTD(**pY**)VATR | 99% | 48.41 | 1/2 | 2/2 | 2/2 |
| AT2G43790 | ATMPK6, MPK6, MAPK6, ATMAPK6 \| MAP kinase 6 | VTSESDF(oxM)TE(**pY**)VVTR | 100%, 100% | 96.41 | 2/2 | 2/2 | 2/2 |
| AT3G45640 | ATMPK3, MPK3, ATMAPK3 \| MAP kinase 3 | PTSENDF(oxM)TE(**pY**)VVTR | 100%, 100% | 58.72 | 2/2 | 1/2 | 2/2 |
| AT1G10210 | ATMPK1, MPK1 \| MAP kinase 1 | GQF(oxM)TE(**pY**)VVTR | 100%, 100% | 46.68 | 2/2 | 1/2 | 2/2 |

### Sequence analysis of D-group MPKs

*Arabidopsis thaliana* encodes 20 mitogen activated protein kinases (MPKs) that phylogenetically form four distinct clusters designated Groups A-D (fig. S1). Unlike Groups A, B and C, the D-Group MPKs have a 60-80 amino acid C-terminal extension (*11*). Alignment of their activation loop, or T-loops, reveals that the D-Group enzymes are also distinct from the A, B and C group MPKs in this region (fig. S2). MPKs are thought to require dual phosphorylation on both the T and Y (TXY motif) of the T-loop for full kinase activation (*12*). Uniquely, the D-group enzymes have an aspartate (D) in the activation loop motif (TDY), while the other groups have glutamate (E) (TEY). Just N-terminal to this motif, but still within the activation loop (defined by DFGX_19-25_ APE), the D-group MPKs have an insert making this surface exposed loop longer, and uniquely each insert has a conserved putative protein phosphatase one (PP1) docking site (defined as RVXF (*2, 4*)), which is not seen in the A, B and C groups (fig. S2 and S3). The MAPK enzymes, including the Arabidopsis A, B and C groups have a common docking (CD) domain, that has been characterized as a common binding site for the activating MAPK kinase (MAPKK), the inactivating MAPK phosphatase, and MAPK substrates via their KIM (Kinase Interacting Motif) sequence (*11, 13–15*). Further adding to the uniqueness of the plant D-group enzymes, unlike the A, B, and C-group MPKs, the CD domain is absent in D-group MPKs (*11, 14*). This absence is consistent with the notion that D-group MPKs do not appear to be phosphorylated by an upstream kinase and auto-phosphorylate to activate (*16*). Consequently, we predict D-Group MPKs have evolved a mechanism for dephosphorylation and inactivation independent of docking known MAPK phosphatases via a CD domain (*11, 13*). Together, this suggests that D-group MPKs autophosphorylate to activate and are likely dephosphosphorylated by a unique mechanism compared to other MPKs. Consistent with this, there is no obvious KIM motif in AtRLPH2 (*7*).

### AtRLPH2 preferentially interacts with D-group MPKs

With phospho-proteomics having identified the D-group MPKs as putative substrates of AtRLPH2, we focused on MPK9 as a representative of the D-group MPKs and compared it to the A-group enzyme, MPK3. We began by exploring the affinity of RLPH2 for MPK9 and MPK3 with overlay dot blots with full length purified and activated/phosphorylated MPKs. AtRLPH2 readily binds the TDY containing MPK9 but shows reduced association if the activation loop TDY aspartate is mutated to a glutamate (TEY) (Fig. 2c). AtRLPH2 displayed almost no binding to the TEY containing A-group MPK, MPK3 and changing the TEY motif glutamate to aspartate (TDY) did not increase affinity (Fig. 2d). This suggests that although the TDY motif plays a role in binding/substrate specificity, another feature of the D-group enzymes provides docking specificity. Notably, only the D-Group enzymes have a long C-terminal tail (*16*).

**Figure 2.**
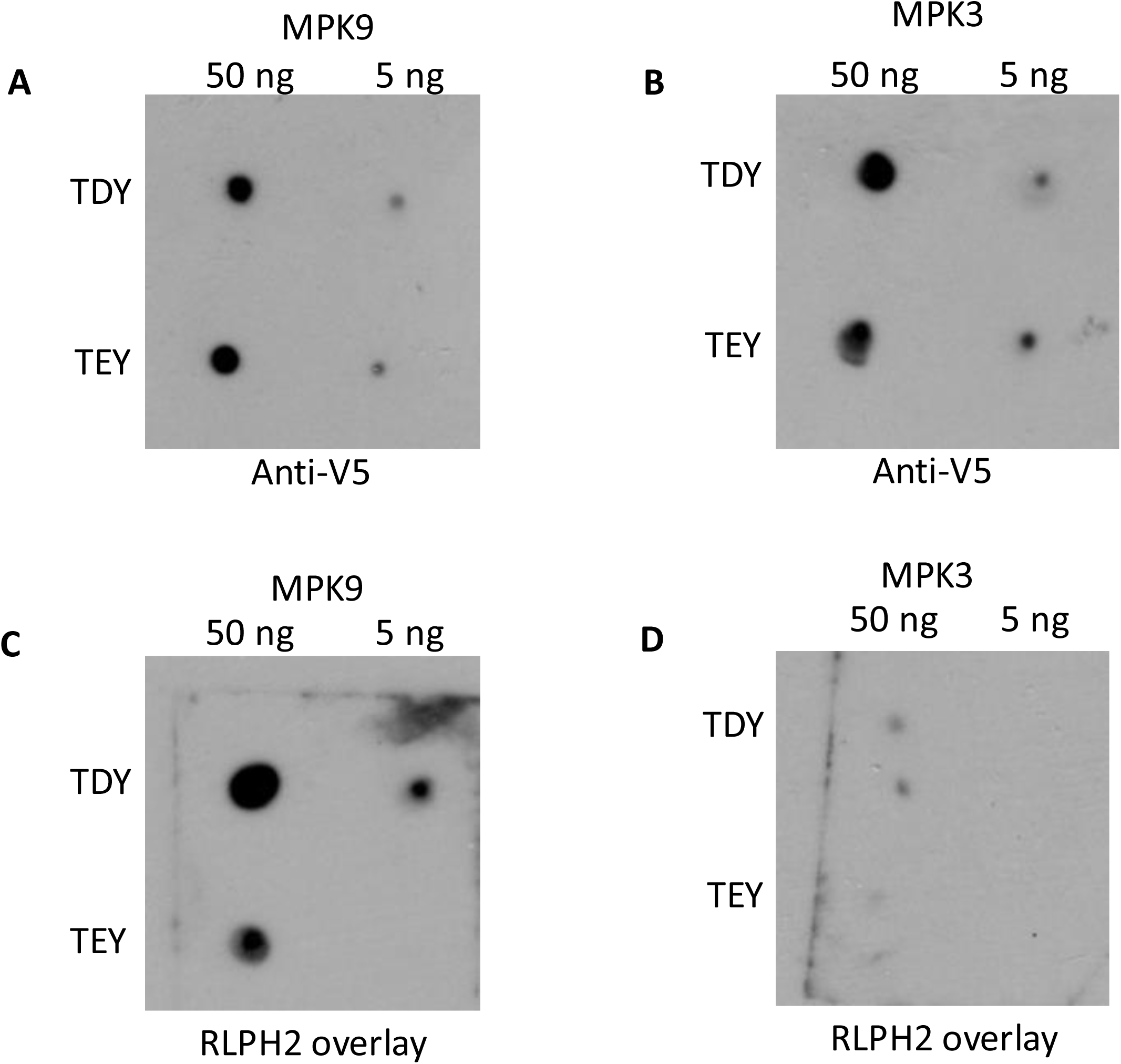
AtRLPH2 preferentially interacts with a D-group (MPK9) MPKs. Purified MPK9 (TDY) and MPK3 (TEY) proteins, along with activation loop mutated versions MPK9 (TEY) and MPK3 (TDY) were spotted to a nitrocellulose membrane for overlay analysis with purified AtRLPH2 (1.5µg/mL). Spotted MPK9 and MPK3 were probed with anti-V5 for equal loading confirmation (**A** & **B**) or overlaid with AtRLPH2. Phosphatase binding was then detected with anti-RLPH2 antibody (**C** & **D**). All experimentation was performed in parallel to ensure comparability.

### AtRLPH2 preferentially dephosphorylates D-group MPKs

Next, we examined the ability of AtRLPH2 to dephosphorylate phosphopeptides derived from the activation loop of MPK9 (Group D MPK) and MPK3 (Group A MPK). Previously we demonstrated that AtRLPH2 has an enhanced V_max_ and decreased K_m_ for an ERK1/2 derived activation loop dually phosphorylated peptide substrate (pTxpY), compared to the same peptide solely phosphorylated on tyrosine (pY) (*7*). Using dually phosphorylated peptides derived from the activation loops of MPK3 and MPK9, AtRLPH2 displayed 19-fold higher activity for the MPK9 substrate versus MPK3 (Fig. 3a). One of the key differences between the MPK9 (GLARVSFNDAPSAIFWpTDpYVATR) and MPK3 (DFGLARPTSENDFMpTEpYVVTR) activation loop peptides is the D versus E in the TXY motif (see fig. S2). When we altered the MPK9 peptide solely at the TXY motif by converting the aspartate (D) to glutamate (E) (GLARVSFNDAPSAIFWpTEpYVATR), we observed a remarkable ∼12-fold decrease in activity (Fig. 3a). This indicates that much of AtRLPH2’s substrate specificity comes from this single amino acid change and supports the idea that AtRLPH2 specifically targets D-group MPKs *in vivo*, as suggested from phospho-proteomic dataset (Table 1).

**Figure 3.**
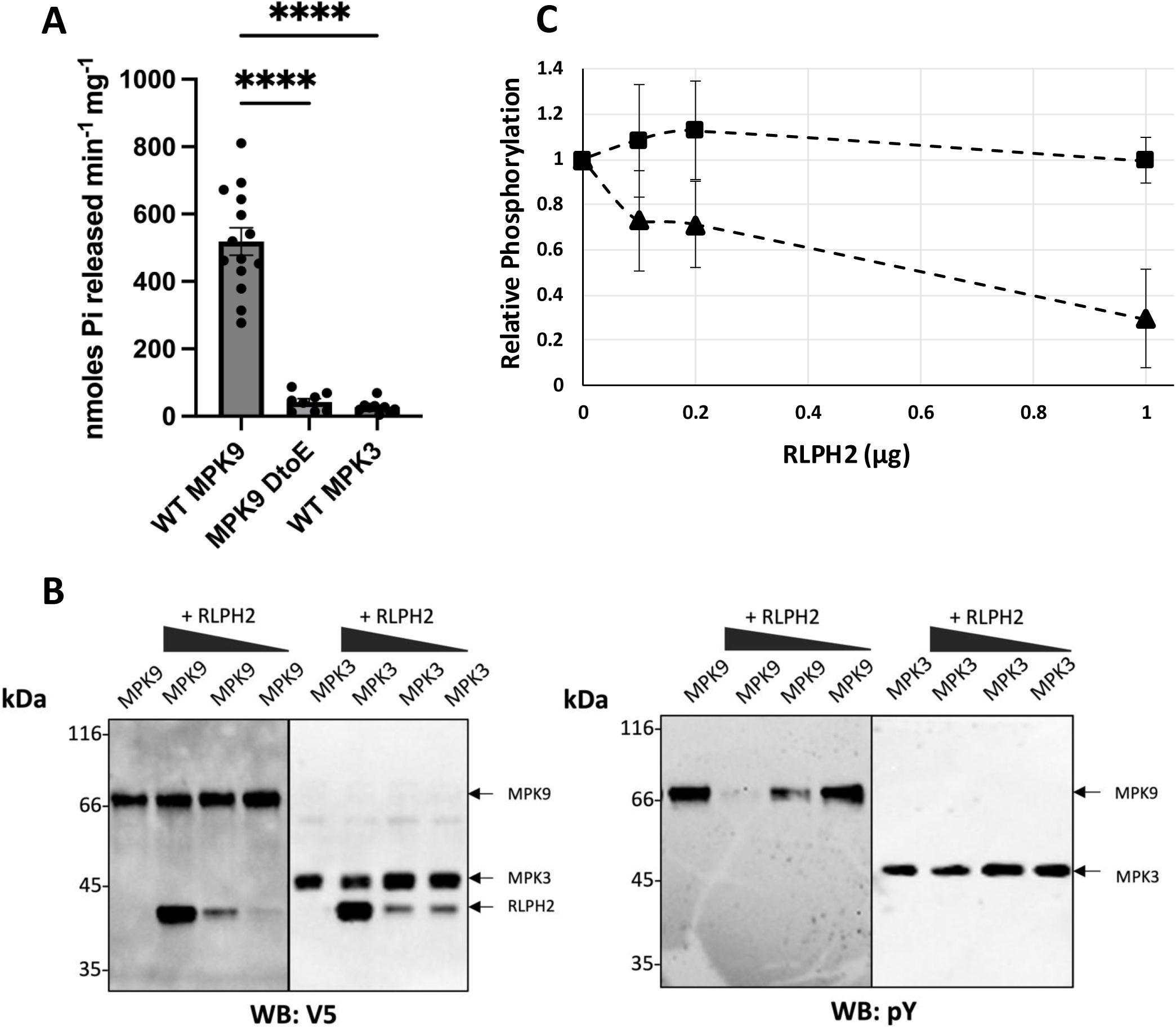
AtRLPH2 preferentially dephosphorylates D-group (MPK9) MPKs. Purified AtRLPH2 was used for in vitro phosphopeptide assays (**A**) based on the dually phosphorylated activation loops of MPK9 (GLARVSFNDAPSAIFWpTDpYVATR), mutated MPK9 (GLARVSFNDAPSAIFWpT**E**pYVATR), and MPK3 (DFGLARPTSENDFMpT**E**pYVVTR) where pT and pY are phospho-threonine and phospho-tyrosine respectively. Shown is the mean where each assay was done in duplicate and repeated either 15 times for TDY MPK9 or 8 times for TEY MPK and MPK3. Comparison of TDY MPK9 to TEY MPK9, and MPK3 using a statistical one-way ANOVA test was conducted and showed a significant difference between D MPK9 and the other two assays. P value for the difference was <0.0001 using a one-way ANOVA. (**B**) varying amounts of RLPH2 was used to dephosphorylate recombinant MPK9 or MPK3 in vitro that was dually phosphorylated prior to the assay. One blot (left) was used to show equal loading using the V5 tag (WB: V5), while the other (right) monitored phosphorylation status of the pY after the assay (WB: pY). Representative blots are shown while in (**C**) the relative phosphorylation from three sets of blots is shown (MPK9, triangles; MPK3, squares). Here the signal intensity of the pY signal is plotted relative to the total protein (V5) signal. Mass standards are shown in kDa.

Next, we examined the ability of AtRLPH2 to dephosphorylate full-length recombinant MPK9 or MPK3 *in vitro*. MPK3 proteins were phosphorylated by the upstream kinase kinase (CKK4DD (*17*)) and MPK9 proteins were allowed to auto-phosphorylate resulting in active, dual phosphorylated (pTxpY) (*18*) MPKs (fig. S4). As shown in Fig. 3b and 3c, AtRLPH2 readily dephosphorylated the pY of MPK9, but did not dephosphorylate the pY of MPK3. Additional assays using WT MPK3 and MPK9 or with switched TDY and TEY motifs, demonstrate that altering D for E in MPK9 drastically reduces the ability of AtRLPH2 to dephosphorylate MPK9 (fig. S4b), while the reverse change on MPK3 (E to D) did not increase dephosphorylation of MPK3. We also found that AtRLPH2 has little to no ability to dephosphorylate the pT of the TXY motif (fig. S4c) (*7*), further supporting the notion that AtRLPH2 is a D-Group MPK tyrosine specific phosphatase.

### Loss of AtRLPH2 increases D-Group MPK T-loop phosphorylation in vivo

We next generated a phospho-specific antibody targeting the activation loop of MPK9 (SAIFWpTDpYVATR) to determine if we could observe increased phosphorylation of MPK9 in a AtRLPH2 knockout line. Notably, the activation loop of D-Group MPKs are very similar (fig. S5) and so we expected that the MPK9 activation loop phospho-specific antibody would recognize these enzymes as well. A, B, and C-group MPKs all have predicted masses of ∼42-45 kDa, while D-group enzymes have predicted masses of ∼55-69 kDa. The antibodies were shown to be phospho-specific (fig. S5b). By comparing WT to *atrlph2* plant extracts we observed proteins corresponding to the expected size to be D-Group enzymes displaying increased phosphorylation in the absence of AtRLPH2 (Fig. 4).

**Figure 4.**
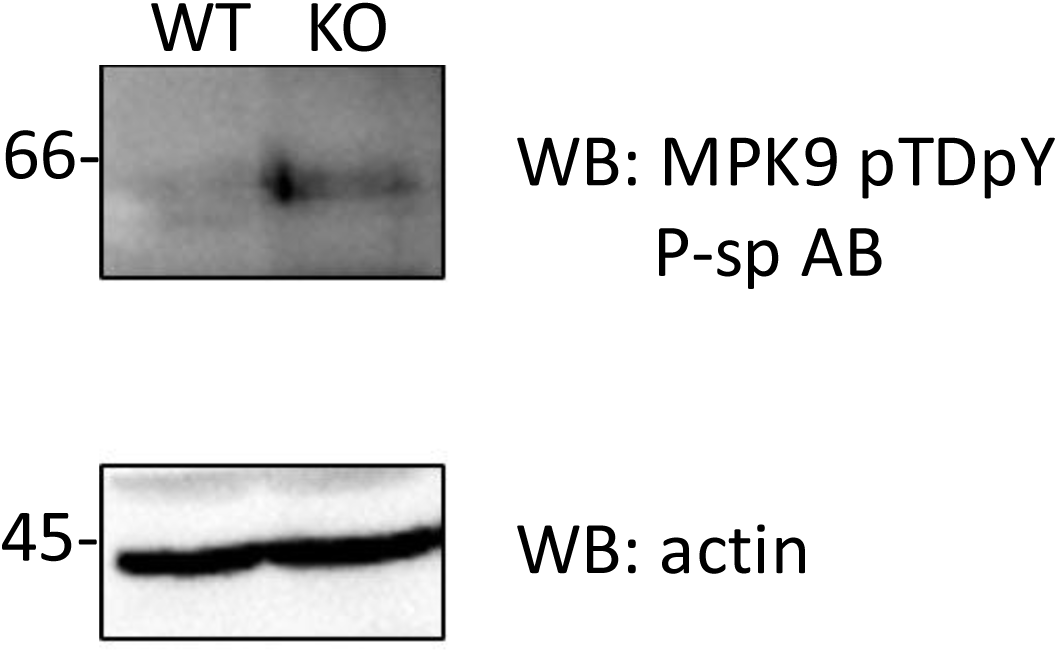
Endogenous D-group MPKs display increased TDY phosphorylation in the absence of protein phosphatase AtRLPH2. Clarified rosette extracts from WT and *rlph2* plants (KO) were run on SDS-PAGE, blotted and probed with a phospho-specific (P-sp) antibody that recognizes the pTDpY sites of D-group MPKs (top panel). The same membrane was cut at ∼50 kDa and probed for actin to show equal protein loading (lower panel). Mass standards are shown in kDa.

### D-group MPKs bind protein phosphatase one (PP1)

Our data indicates that AtRLPH2 specifically binds D-Group MPKs and selectively dephosphorylates the activation loop pY of these enzymes. This is facilitated by recruiting the MPK via docking the pT in the AtRLPH2 basic pocket adjacent to the active site (*7*). However, complete inactivation of MPKs typically requires the dephosphorylation of both TXY motif residues, which AtRLPH2 does not appear to do. Interestingly, as discussed above, the D-group MPKs possess an insert in their activation loop (fig. S2 and S3) containing a potential docking site for a serine/threonine specific PP1 phosphatase (PP1 isoforms are named TOPP1-TOPP9 in Arabidopsis (*4*)). In eukaryotes, the RVXF motif functions as a PP1 recruitment motif, allowing the phosphatase to be regulated by a series of protein interactors that direct its function in the cell by guiding PP1 catalytic subunits to their corresponding substrates (*2, 4, 19*). A subset of PP1 interactors also have additional binding residues C-terminal of the RVXF motif (RVxF_5-8_ΦΦ_8-9_R, where Φ is a hydrophobic residue) (*20*), a feature also present in D-group MPKs (fig. S3). This, combined with the fact that activation loops are on the surface of protein kinases, supports the hypothesis that these D-Group MPKs also interact with PP1 through the RVXF motif to potentially drive dephosphorylation of the pT residue of the TDY motif, or to target other phosphorylation sites in other proteins, or even the abundant phospho-sites of the C-termini of D-group MPKs (*18, 21*). Importantly, the RVXF motif insert found in D-Group MPK activation loops is not found in the Group A, B or C MPKs (fig. S2). Further analyses of plant and other eukaryotic protein kinases found no evidence for any other protein kinase activation loops, in any organism, to have this PP1 docking insert, making this another unique attribute of plant D-Group MPKs. Given the conservation of the novel TDY activation loop and the putative PP1 binding site across plant D-group MPKs [see additional file 4 in (*11*) and our observations], this evidence suggests that docking PP1 is fundamental to the function of D-Group enzymes.

Given these observations, we proceeded to explore whether PP1 indeed binds to D-group MPKs. To do this, we first investigated a MPK9 model generated by AlphaFold2. As predicted, the RVXF motif is surface exposed (fig. S6). We then expressed and purified each Arabidopsis PP1 (TOPP1-TOPP9 (*22*)) using the affinity matrix microcystin-Sepharose (*23*) and performed overlays using purified MPK9 where the putative PP1 binding RVXF motif was maintained or mutated to RASA (non-PP1 binding (*4, 19*)) in each TOPP. This demonstrated that each PP1 isoform can bind MPK9, and this is dependent on the key hydrophobic residues of the RVXF sequence (Fig. 5a; note, Arabidopsis PP1 isoform 6 does not bind to microcystin and thus did not purify on the affinity matrix (*22*)). Next, we used Arabidopsis PP1 isoform 5 as a representative to perform an *in vitro* pulldown between purified MPK9 and PP1. As predicted, this association was dependent upon the RVXF motif. (Fig. 5b).

**Figure 5.**
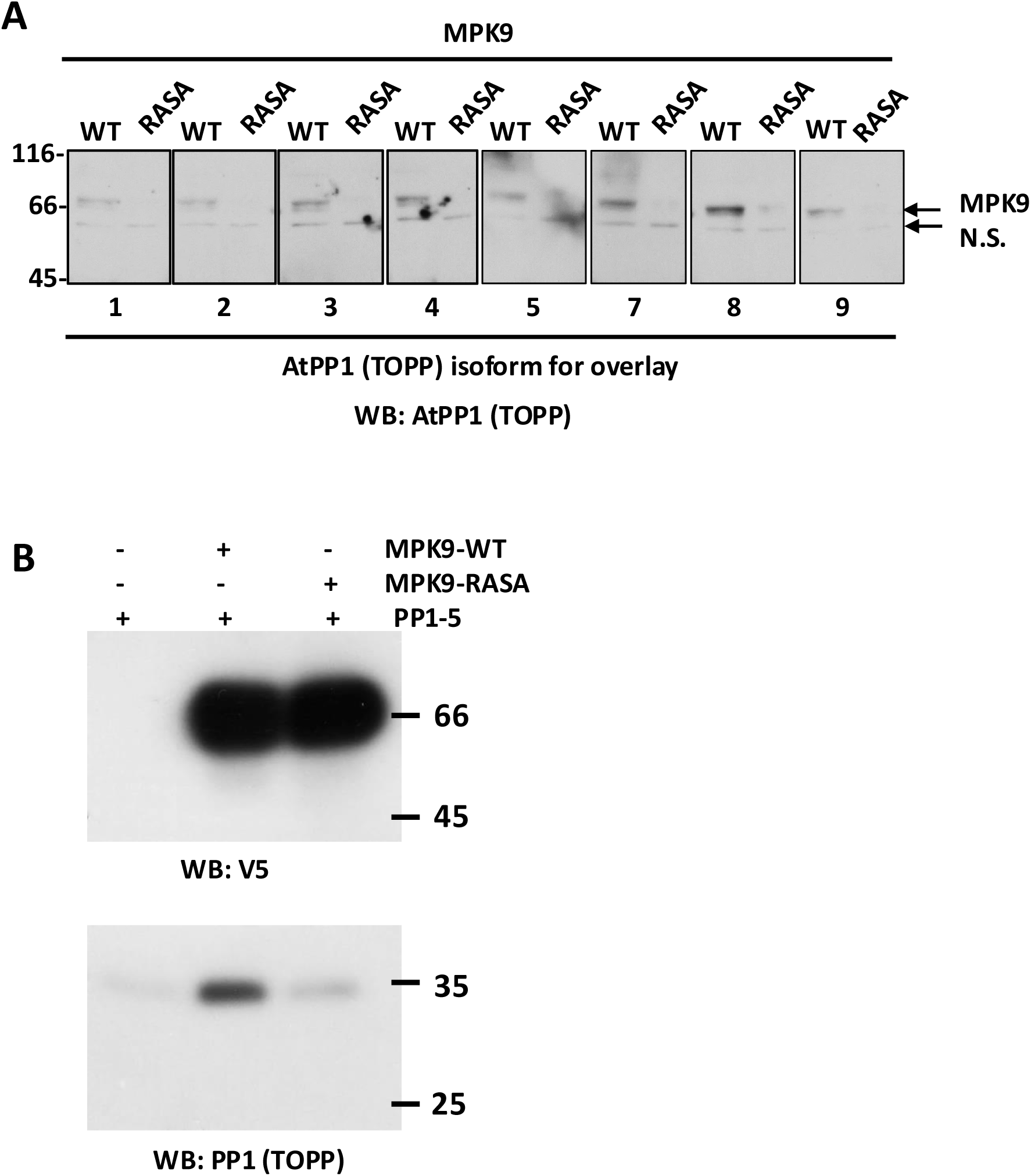
All PP1 (TOPP) isoforms bind the D-group MPK, MPK9. (**A**) Purified WT MPK9 or MPK9 with mutated putative PP1 binding RVSF motif (RASA) were incubated (overlaid) with purified recombinant Arabidopsis PP1 isoforms (TOPPs) 1 through 9 (except PP1isoform 6) as indicated below each box. Binding of PP1 to MPK9 was visualized with an anti-PP1(TOPP) antibody. N.S. is a non-specific band detected by the PP1 antibody in all samples. (**B**) Pulldown assays were performed with WT and RASA-mutated MPK9 (5 µg) with PP1 isoform 5 (1µg, TOPP5) as the representative PP1. MPK9 was detected with a V5 epitope antibody and PP1with a pan-TOPP1 isoform antibody (*22*). Experimentation with PP1 isoform 6 (TOPP6) was not performed as it does not bind and purify on microcystin-Sepharose. Mass standards are shown in kDa.

### AtRLPH2 represses seed dormancy release, and affects seed sensitivity to gibberellic acid synthesis inhibitors and abscisic acid

Few D-Group MPKs have defined functions, with one exception being MPK8, which has been characterized in promoting seed germination (*10*). Therefore, to address a possible role of AtRLPH2 in seed germination, freshly harvested AtRLPH2-OE (Columbia ecotype) and *atrlph2* (Nössen ecotype) seeds were germinated together with the corresponding WT lines at 15°C (Fig. 6a). In these conditions, the germination of all seed lines was over 89% after 7 days. The *atrlph2-1* and *atrlph2-2* mutant lines exhibited accelerated germination compared to WT Nössen. To compare the dormancy level of the different seed lines at harvest, germination was also performed at 25°C (Fig. 6a). In these conditions, *atrlph2* mutant seeds presented significantly higher germination than WT Nössen seeds (5.6%), with *atrlph2-2* germinating better than *atrplh2-1* (72 and 20%, respectively). Conversely, the AtRLPH2-overexpressing line presented a lower germination rate compared with WT Col-0 seeds (41 and 70.4% respectively). These data indicate that AtRLPH2 is a positive regulator of seed dormancy.

**Figure 6.**
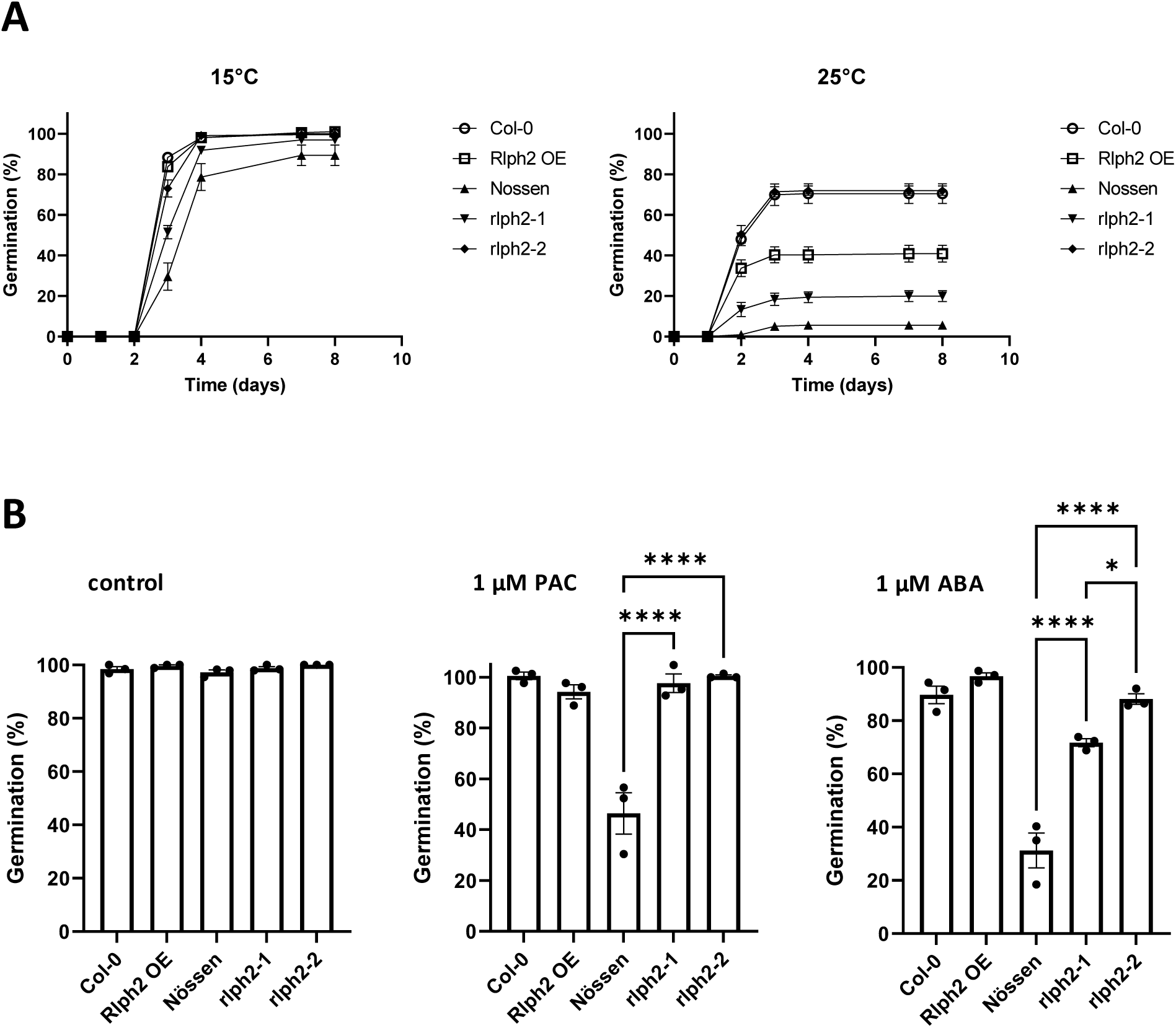
Germination phenotypes of RLPH2 knockout and overexpressing seed lines. (A) Seeds from WT Nössen (solid triangle), *rlph2-1* (solid triangle), *rlph2-2* (solid diamond), Col-0 (open circle) and 35S::RLPH2-cTAP (RLPH2 OE) (open square) were incubated at 15°C and 25°C in darkness for 8d. Results represent the mean germination percentage ± SE of four replicates of 50 seeds. (B) Assessment of ABA and paclobutrazol (PAC) sensitivity of non-dormant seeds. Here, Nössen, *rlph2-1*, *rlph2-2*, Col-0 and RLPH2 OE seeds were stratified 4d at 4°C on MS agar medium (control) or in the presence of 1 µM ABA or 1 µM PAC. The germination rate was scored after 7d on light at 21°C. Results represent the mean germination percentage ± SE of three replicates of 50 seeds. Asterisks indicate statistical differences between Nössen and *rlph2* mutants or between Col-0 and RLPH2 OE, as determined by One-way ANOVA with post hoc Tukey’s test (*P* < 0.05).

Next, the sensitivity of the different seed lines to ABA and to the GA synthesis inhibitor paclobutrazol (PAC) was compared after 7 days of germination. Dormancy was released by stratification and all seeds fully germinated in the absence of treatment (Fig. 6b). In contrast, in the presence of 1 µM ABA, *atrlph2-1* and *atrlph2-2* seeds exhibited a higher germination rate than WT Nössen seeds (71.7-88.1% and 31.2%, respectively). At higher ABA concentrations, the germination of the three lines was almost fully inhibited (fig. S7). On the other hand, *atrlph2* mutants were less affected by 1 µM PAC compared to WT Nössen seeds (97.6-100% and 46.4% germination, respectively) (Fig. 6b). This difference was also observed at higher PAC concentrations (fig. S8). No difference of sensitivity to ABA and PAC was observed between Col-0 and AtRLPH2-overexpressing line seeds at any concentration (Fig 6b, fig. S7 and S8). These data suggest that the function of AtRLPH2 during seed germination involves ABA and GA pathways.

## Discussion

MPKs are ubiquitous eukaryotic signaling proteins. In all cases they are activated by phosphorylation of their activation or T-loops within a characteristic TXY motif (*15*). This event, as in all protein kinases, re-shapes the active site repositioning key catalytic residues (*24*). In plants, the MPK family has expanded compared to yeast and humans (*11, 14*) and in Arabidopsis includes 20 members (fig. S1). These members form four groups based on sequence, with the D-Group having unique sequence additions. Then importance of the D-Group MPKs was emphasized in a recent analysis of 40 plant genomes, including eukaryotic algae, bryophytes, monocots and dicots. Here, they found evidence that some species have lost one of the A, B or C groups, but each species maintained multiple D-Group TDY enzymes (*25*).

In this study, we provide evidence that the plant tyrosine phosphatase AtRLPH2 specifically targets and dephosphorylates the pY residue of the TDY motif in D-Group MPKs. These enzymes are unique in possessing a TDY activation loop motif, a PP1 docking SLiM in the same loop, and a C-terminal extension with multiple mapped phosphorylation sites (*18*). The specificity of AtRLPH2 toward the D-Group MPKs seem to involve two components: 1) In an overlay assay AtRLPH2 readily binds TDY-MPK9 and this is marginally reduced in TEY-MPK9. 2) AtRLPH2 does not bind the A-Group MPK3 regardless of the T-loop motif being TEY or TDY, suggesting that this interaction is driven by some feature outside of the TXY motif-perhaps the C-terminal tails of the D-Group MPKs. Indeed, alignment of the D-Group MPK C-terminal tails reveals a conserved core sequence (fig. S9). In dephosphorylation assays using phosphopeptides, changing the D to an E in the MPK9 substrate peptide drastically reduces activity while the TEY-MPK3 phosphopeptide is simply a poor AtRLPH2 substrate, suggesting substrate sequence specificity is conferred by the active site. This is supported by dephosphorylation assays of the recombinant enzymes leading to a model where AtRLPH2 achieves D-group specificity by associating with some structural feature unique to D-Group MPKs, then a second layer of specificity is driven by the active site itself, selecting for TDY motifs.

Classically, it is been believed that full activation and inactivation of MAPKs require dual TXY phosphorylation and dephosphorylation, respectively; however, the mono-phosphorylated versions have been detected in human cells (*26*) and may serve as reservoirs of MAPK that can be rapidly converted to fully activated in cells (*26*). Most MAPKs are dephosphorylated by specific MAPK phosphatases which target both the T and Y of the loop, yet human dual specificity phosphatase VHR/DUSP3, like AtRLPH2, recruits specific MAPKs via a pT binding basic pocket to specifically dephosphorylate solely the pY (*9*). The mono-phosphorylated product could then be potentially acted upon by specific PP2C (*12*) or PP2A (*12, 15, 27*) serine/threonine phosphatases. The activity of the mono-phosphorylated MAPKs seems to vary depending on the enzyme, based on the few detailed studies. For instance, the protein kinase activity of human p38/pY and p38/pT are about 1000- and 10-fold lower than that of the dual phosphorylated p38/pTXpY, respectively, while mono-phosphorylated (pY or pT) ERK2 displays activities intermediate of full and non-phosphorylated enzyme (*12*). This led to the speculation that mono-phosphorylated versions could have unique substrate specificity/biological roles or act as a reservoir to rapidly activate to fully phosphorylated enzyme (*12*). Whether plant D-Group MPKs only phosphorylated on threonine (pT) display these properties is unknown and is worthy of further future investigation.

Observationally, we identified a putative PP1 binding SLiM in all D-Group MPK activation loops (a feature absent in A, B and C Group MPKs) and consistent with being surface exposed we show this sequence does bind serine/threonine phosphatase 1 (PP1). Interestingly, through the use of the phosphatase inhibitor okadaic acid, the activation loop of MPK9 was shown to be regulated *in vivo* by an okadaic acid sensitive phosphatase (PP2A, PP1, PP4-PP6) (*18*) and we previously showed that AtRLPH2 activity is not affected by this inhibitor *in vitro* (*8*). We therefore speculate that the recruited PP1 may dephosphorylate the pT of the MPK T-loop itself, target the C-terminal tail phosphorylation sites (*18, 21*), or could simply target other proteins linked to D-Group MPK function. It is also worth noting that five of the eight D-Group activation loop RVXF motifs have a serine in the X-position (RVSF; fig. S2) and phosphorylation of this site would block PP1 binding (*28*), as has been shown for RVSF motifs in other PP1 interactors, potentially bringing another level of regulation to D-group enzyme regulatory events.

Finally, we began exploring AtRLPH2 function *in planta*. Looking for the already known functions of the D-Group MPKs, we noticed their functions have not been well studied, with one exception being MPK8 which has previously been linked to the promotion of seed dormancy release. If AtRLPH2 regulates D-Group MPKs we reasoned that loss of AtRLPH2 could keep MPK8 phosphorylated and therefore active, promoting germination. The germination capacity of *atrlph2-1* and *atrlph2-2* revealed accelerated germination rates, therefore AtRLPH2 acts as a repressor of germination, perhaps through keeping MPK8 dephosphorylated on its T-loop tyrosine. Further, with MPK8 involvement in dormancy release shown to be gibberellic acid (GA) dependent, we investigated the involvement of AtRLPH2 in GA metabolism. Contrary to *mpk8* mutants, both *atrlph2* mutants were significantly less sensitive to the GA biosynthesis inhibitor paclobutrazol (PAC) suggesting AtRLPH2 is a negative regulator of seed germination by either negatively regulating GA synthesis or signaling, or positively regulating GA catabolism. This could be via MPK8 inactivation by dephosphorylation, leading to reduced or no phosphorylation of the MPK8 downstream target, TCP14 (*10*). With seed germination orchestrated by a balance of gibberellic acid*/*abscisic acid (ABA) signaling, ABA sensitivity of *atrlph2* mutants was also assessed. Both lines were significantly less sensitive to 1 µM ABA than the WT Nössen plants. This suggests that AtRLPH2 is also involved in negatively regulating seed germination by being a positive regulator of ABA biosynthesis/signaling or a negative regulator of ABA catabolism.

We have revealed the D-Group MPKs as targets of the novel plant tyrosine phosphatase RLPH2 and uncovered that its substrate specificity relies in part on a single amino acid substitution (D or E in the TXY motif) in substrates. We also demonstrate the unique ability of D-Group MPKs to recruit PP1 via a conserved RVXF SLiM. Finally, we find that AtRLPH2 is involved in inhibition of seed germination in a GA and ABA dependent manner.

## Materials and Methods

### Phospho-proteomics sample preparation

Wild type (WT) and two independent *rlph2* (*rlph2-1* and *rlph2-2*) mutant allele containing *Arabidopsis thaliana* plant lines were grown as described in (*8*). Rosette leaves were harvested, flash frozen in liquid nitrogen and stored at −80°C until day of use. Protein extraction and digestion was performed by Filter Aided Sample Preparation (FASP) as previously described without deviation (*29*).

Phosphorylated peptides were then enriched using TiO_2_ beads, at a concentration of 10 mg/mL as previously described (*30*). Eluate phosphopeptides were then acidified with formic acid to a concentration of 0.5% (v/v) and dried with speed vacuum (ThermoFisher). Dried phosphopeptides were then re-suspended in 800 µL ice-cold immunoprecipitation (IP) buffer (50 mM HEPES-NaOH pH 7.4, 50 mM NaCl), pH adjusted to 7.5 and incubated end-over-end, overnight at 4°C with 50 µL of anti-phospho-tyrosine antibody agarose beads (Santa Cruz, sc-7020) and 15 µL of anti-phospho-tyrosine antibody Sepharose beads (p-Tyr-100, Cell Signaling, #7902) pre-equilibrated in IP buffer. Beads were pelleted and washed 3 times for 2 min at RT with 1 mL IP buffer followed by 3 washes with ddH_2_O. Enriched tyrosine phosphorylated peptides were then eluted with 3 x 200 µL of an 80% (v/v) ACN /2% (v/v) formic acid (FA)solution. Eluates were pooled and dried by speed vacuum (ThermoFisher).

### Mass Spectrometry Analysis

All dried phosphopeptide samples were dissolved in a 3% ACN (v/v) / 0.1% (v/v) TFA solution and desalted using ZipTip C18 pipette tips (Millipore; ZTC18S960) as previously described (*29*). Phosphopeptides were then dried and dissolved in a solution of 3.0% ACN (v/v/) / 0.1% FA (v/v) solution prior to mass spectrometry analysis. Dissolved samples were then injected using an Easy-nLC 1000 system (ThermoFisher) and separated on a 50 cm EasySpray nano-LC column (ES803). The column was equilibrated with 100% solvent A (0.1% FA in water). Peptides were eluted using the following gradient of solvent B (0.1% FA in ACN): 0-50 min; 0-25% B, 50-60 min; 25-32% B, 60-70 min; 32-98% B at a flow rate of 0.3 µl/min. High accuracy mass spectra were acquired using an Orbitrap Fusion mass spectrometer (ThermoFisher) that was operated in data dependent acquisition mode. All precursor signals were recorded in the Orbitrap using quadrupole transmission in the mass range of 300-1500 m/z. Spectra were recorded with a resolution of 120 000 at 190 m/z, a target value of 4E5, and the maximum cycle time was set to 3 seconds. Data dependent MS/MS were recorded in the linear ion trap using quadrupole isolation with a window of 2 Da and HCD fragmentation with 30% fragmentation energy. The ion trap was operated in rapid scan mode with a target value of 3E4 and a maximum injection time of 100 ms. Precursor signals were selected for fragmentation with a charge state from +2 to +7 and a signal intensity of at least 1E4. A dynamic exclusion list was used for 30 seconds and maximum parallelizing ion injections was activated. After data collection peak lists were extracted from the instrument raw files using Proteome Discoverer (version 1.4/2.1) and exported to mgf format for subsequent database analysis. All MS/MS samples were analyzed using Mascot (Matrix Science, version 2.5.3).

Tandem MS spectra were then searched with Mascot 2.5 ((http://www.matrixscience.com; Matrix Science, London, UK) against The Arabidopsis Information Resource (TAIR10) protein database using a concatenated decoy database supplemented with contaminants. Mascot search parameters included: trypsin specificity, peptide mass tolerance of 10 ppm, MS/MS tolerance of 0.6 Da, charge state of +2, +3 or +4 and fixed cysteine carbamidomethylation. Variable modifications included: serine, threonine, tyrosine phosphorylation and methionine oxidation with 1 missed cleavage. Mascot identifications were then imported into Scaffold Q+S ((http://www.proteomesoftware.com) and filtered for a protein and peptide false discovery rate (FDR) ≤ 0.01 (table S2). Scaffold data were then assessed for pY phosphoprotein identifications unique to *rlph2-1* and *rlph2-2* (table S2).

### Cloning, expression and purification of enzymes

AtRLPH2 was expressed and purified as in (*8*). Synthesized MPK9 (At3g18040) was codon optimized for bacterial expression (GenScript). AtMPK9 was then cloned into bacterial expression vector pDEST42 (Invitrogen) for expression using BL21 (DE3) CodonPlus-RIL *E. coli.* The resulting MPK9-V5-His6 construct was then produced in BL21 (DE3) CodonPlus-RIL *E.coli* as follows: Bacteria were grown at 37°C until an OD_600_ of 0.4-0.6, followed by induction with 0.1 mM IPTG for 7 hr at 37°C. Induced cells were centrifuged at 3300xg for 20 min, with the resulting cellular pellet re-suspended in 15 mL of resuspension buffer (50 mM HEPES pH 7.5, 150 mM NaCl, 5% v/v glycerol, 10 mM imidazole) plus 1 mM benzamidine and 1 mM PMSF. Bacterial cells were then lysed by French press three times at 1000psi, followed by centrifugation at 50,000xg for 45 min. The resulting supernatant was then collected and filtered through miracloth before a 1 hr incubation with 1 mL of Ni-NTA agarose (Qiagen). Matrix was washed with buffer A (50 mM HEPES pH 7.5, 1 M NaCl, 5% glycerol, 10 mM imidazole, 0.5% Tween-20) followed buffer B (50 mM HEPES pH 7.5, 150 mM NaCl, 5% glycerol, 10 mM imidazole) before being eluted with 400 mM imidazole in 50 mM HEPES pH 7.5, 150 mM NaCl, 5% glycerol. Purified protein was then concentrated in an Amicon 30 kDa centricon (Millipore, UFC9030), and buffer was exchanged for wash buffer B. Protein concentration was determined by Bradford using BSA as standard and enzyme was stored at −80°C until use.

MPK3 (At3g45640) in pDEST42 (MPK3-V5-His6) was purified, concentrated, and stored like MPK9, except bacteria were only induced for 4 hrs. A constitutively active CAMKK4-GST for MPK3 phosphorylation and activation was expressed in a BL21 (DE3) CodonPlus-RIL *E. coli* strain (*17*). Cells were grown until an OD_600_ of 0.4-0.6 was reached, at which point 0.1 mM IPTG was added to induce expression for 6 hr at 30°C. Cells were centrifuged at 3300g for 20 min at 4°C, after which the pellet was resuspended in 15 mL of resuspension buffer (50 mM HEPES pH 7.5, 150 mM NaCl, 3 mM DTT, 0.05% NP-40, 2 µg/mL leupeptin, 5 µg/mL pepstatin) and lysed by French press, followed by centrifugation at 50,000g for 45 min. The supernatant was collected and filtered through miracloth before a 1 hr incubation with 1 mL of glutathione-Sepharose 4B matrix (Cytiva). Matrix was washed with 300 CV of wash buffer C (50 mM HEPES pH 7.5, 750 mM NaCl, 1 mM DTT, 0.1% NP-40) followed by an additional 100 mL of wash buffer D (50 mM HEPES pH 7.5, 150 mM NaCl, 1 mM DTT) before elution with 20 mM reduced glutathione in 50 mM HEPES pH 7.5, 150 mM NaCl, and 1 mM DTT. Protein was concentrated as for MPK9.

The Arabidopsis protein phosphatase one (PP1) enzymes (TOPP 1-TOPP9) were purified on microcystin-Sepharose as described (*22*). PP1 isoform 6 (TOPP6) was not purified because it is not microcystin sensitive.

### Site directed mutagenesis

Using the QuikChange Lightning Site-Directed Mutagenesis Kit (Agilent Technologies) mutants of MPK9 and MPK3 were generated by point mutations in their respective activation loops. The MPK9 RASA mutant construct had valine and phenylalanine of the RVSF motif (see **fig. S2**) changed to alanine. The MPK9 TEY mutant construct had the aspartic acid (D) within MPK9s TDY motif switched for a glutamic acid (E). Similarly, the MPK3 TDY mutant construct had its glutamic acid in TEY motif switched for an aspartic acid.

### Phosphorylation of MPK9 and MPK3 activation loops

Purified MPK9, MPK3 and activation loop mutant enzymes were diluted to 500 µL in 25 mM HEPES pH 7.5, 10 mM NaCl, 15 mM MgCl_2_ and 5 mM ATP. MPK9 was allowed to auto-phosphorylate or 500 ng of constitutively active protein kinase CAMKK4(DD) was added to MPK3 and samples incubated at 30°C. After 2 hr Mg-ATP was removed by concentration to 50 µL and dilution back to 500 µL with 50 mM HEPES pH 7.5, 150 mM NaCl, 5% glycerol using a 30 kDa centricon, five times. Protein samples were then stored at −80°C until day of use.

### AtRLPH2 phosphatase assays

For phosphopeptide assays, purified AtRLPH2 (1 µg) was incubated for 30 min at 200 rpm at 30°C in 160 µL of assay buffer (50 mM HEPES pH 7.5, 150 mM NaCl) containing 0.5 mM peptide substrate in a 96-well plate. Peptides were at least 95% pure and purchased from GL Biochem (Shanghai) Ltd. Control assays were performed in parallel with identical conditions but lacking enzyme. The reaction was stopped with the addition of 40 μL malachite green solution and O.D. measured at 630 nm using a plate reader. The activity of AtRLPH2 was measured by comparing absorbance at 630 nm with the standard curve of free Pi prepared in parallel.

For dephosphorylation of purified recombinant and phosphorylated MPK9 and MPK3, MPK (5 µg) was incubated in the absence of, or with 5 µg, 1 µg and 0.5 µg AtRLPH2 in 20 mM HEPES pH 7.5, 10 mM NaCl, 0.1 mM DTT for 1 hr 30 mins at 30°C. Reactions were stopped by the addition of 5X SDS cocktail and heated at 95°C for 10 mins. Dephosphorylation was monitored by blotting with phospho-specific antibodies.

### Western and far-western blots

For MPK proteins dephosphorylated by AtRLPH2, samples were run on a 10% SDS-PAGE and then visualized by immunoblotting with either Horseradish peroxidase (HRP) conjugated rabbit anti-V5 (Immunology Consultants Laboratory, RV5-45P-Z) or anti-pT (Cell Signaling Technologies, 42H4) and anti-pY (Cell Signaling Technologies, PTyr-100) for phospho-status of the MPK protein. V5 antibody was diluted in 5% BSA and 0.1% TBST, and incubated for 2 hrs, and after washing imaged via the conjugated secondary antibody diluted 5000-fold. The anti-pT and anti-pY primary antibodies (1:1000 and 1:5000, respectively) were diluted in 5% BSA and 0.1% TBST and incubated for 2 hrs and after washing probed with anti-mouse secondary antibody diluted 5000-fold. For PP1 far-westerns, WT and RVSF mutated MPKs were run on SDS-PAGE, blotted to nitrocellulose and probed with PP1 isoforms (2µg/mL) for 4 hr in TBS plus 5% milk powder as in Templeton *et al*. (*22*). Antibodies to Arabidopsis PP1 isoforms was generated, affinity-purified and used as in (*22*) to visualize PP1 binding. For dot-blot far-western, purified activated MPKs were dotted onto a nitrocellulose membrane, blocked and probed with anti-V5 (for equal loading) or overlaid with purified AtRLPH2 (1.5µg/mL), and detected with anti-AtRLPH2 antibody as in (*8*).

For activation loop phospho-specific antibody blots samples were generated from Arabidopsis rosettes during the light period by homogenization using a micro pestle in 1.5 mL tubes in 200 µL of 95°C 5X SDS-PAGE loading buffer and 200 µL of ddH_2_O. Samples were then boiled, centrifuged, and the supernatant collected for western blot analysis. The blot was cut at ∼50 kDa and the bottom probed for actin and the top half probed for MPK9 TDY phospho-status. Anti-actin (Agrisera AS13 2640) was diluted 4000-fold in 0.1% TBST plus 5% BSA. Phospho-specific antibodies to the MPK9 activation loop were generated as in Nasa et al. (*28*) with the dually phosphorylated peptide. Briefly 2 mg / injection of the MPK9 T-loop phosphopeptide (SAIFWpTDpYVATR) was dissolved in PBS plus 5 mM of NaPPi, 20 mM NaF and 5 mM of Na_2_OV_4_ and coupled to KLH, quenched with Tris-HCL and dialyzed into the above buffer. Antibodies are characterized in fig. S5.

### PP1-MPK9 pulldown assay

PP1 isoform 5 (1 µg) and MPK9 WT or RASA (5 µg) were incubated together for 1 hr at RT while simultaneously Ni-NTA was incubated with 5 mg/mL BSA to block non-specific interactions. Afterwards PP1 or PP1 with MPK9 WT/RASA was incubated with 0.2 mL Ni-NTA for 1 hr at RT, washed with 50 mM HEPES pH 7.5, 300 mM NaCl Buffer, and 0.1% Tween-20, followed by buffer minus Tween-20, then eluted in 2X SDS cocktail to be analyzed by western blot as described above.

### Germination assays

Germination of freshly harvested seeds was performed in darkness at 15°C or 25°C, as previously described (*31*). For abscisic acid (ABA) or paclobutrazol (PAC) treatments, seeds were sewn on half-strength MS agar medium, supplemented with different concentrations of ABA or PAC, and stratified for 4 d at 4°C. Seeds were then incubated at 21°C under constant light and germination was scored daily for seven days.

## Acknowledgements

We would also like to thank Marcus Samuel for providing the CaMKK4DD and MPK3 clones.

## Author contributions

The study was conceived by AML, RGU and GM. Manuscript writing was a contribution of all authors. Mass spectrometry and analysis was performed by RGU. All biochemical experiments were a combined effort of AML, RT, SM and JJJ. Sequence alignments and AlphaFold structure prediction was done by RT, SM and JJJ. Germination assays were performed by JP and EB.

## Data Availability

All raw data files, mass spectrometry parameters, and Spectronaut search settings have been uploaded to ProteomeXchange (http://www.proteomexchange.org/) via the PRoteomics IDEntification Database (PRIDE; https://www.ebi.ac.uk/pride/). Project Accession: PXD038689.; Password: yqxu8Z7L.

## Supplementary Materials

Fig. S1. Phylogenetic analysis of the 20 MPK proteins of Arabidopsis thaliana.

Fig. S2. Activation loops of Arabidopsis thaliana MPKs.

Fig. S3. D group MPKs have an extended RVXF motif.

Fig. S4. AtRLPH2 solely dephosphorylates pY of MPK9 TDY, but not MPK9 TEY, MPK3 TEY or MPK3 TDY.

Fig. S5. Characterization of MPK9 pTDpY phospho-specific antibodies.

Fig. S6. Predicted Arabidopsis thaliana MPK9 structure.

Fig S7. Percentage germination after 7 days in the presence of ABA.

Fig. S8. Percentage germination in the presence of PAC after 7d.

Fig. S9. Conserved region of C-terminal tails of Arabidopsis D-group MPKs.

Supplemental Table 1. Plant proteins with pTXpY motifs extracted from Plant PTM Viewer (www.psb.ugent.be/webtools/ptm-viewer/) (*32*).

Supplemental Table 2. All pTyr phosphorylated peptides identified by Mascot and Scaffold Q+S.

## Supplemental Figures and Tables

**Figure S1. Phylogenetic analysis of the 20 MPK proteins of Arabidopsis thaliana.**

Groups A, B, C and D are labeled with the corresponding MPKs present in each group. Sequence Viewer 8.0 was used to align the sequences and generate the phylogenetic tree.

**Figure S2. Activation loops of Arabidopsis thaliana MPKs.**

Protein kinase activation loops are defined by the sequences DFG and APE. In MPKs activation is achieved by phosphorylation of a TXY motif (green). Arabidopsis MPKs form 4 phylogenetically distinct groups as indicated (A-D) with only the D-group have an aspartate (D) between the T and Y of the TXY motif, while groups A-C have a E in this position (*11*). The D-group enzymes also possess an insert in the loop that conforms to a protein phosphatase one (PP1) binding RVXF SLiM (blue).

**Figure S3. D group MPKs have an extended RVXF motif.**

The activation loops of the D group MPKs as defined by DFG and APE. Highlighted in blue are the amino acids associated with the extended RVXF motif, which starts with RVXF, then 5-8 amino acids C-terminal two hydrophobic residues (Val, Ile or Phe), and then 8-9 amino acids C-terminal an arginine (R). Each D-group enzyme conforms to this sequence motif further supporting the idea that these protein kinases recruit PP1.

**Figure S4. AtRLPH2 solely dephosphorylates pY of MPK9 TDY, but not MPK9 TEY, MPK3 TEY or MPK3 TDY.**

MPKs were dually phosphorylated in their activation loops (see methods) and treated with or without purified RLPH2. Western blotting was performed to assess the MPK phospho-status with either anti-pY or pT antibodies. (A) Anti-V5 immunoblot demonstrating equal loading. (B & C) Anti-phosphotyrosine and anti-phosphothreonine immunoblots, respectively.

**Figure S5. Characterization of MPK9 pTDpY phospho-specific antibodies.**

(A) Alignment of D-group MPK activation loops highlighting the sequence relationship (boxed green) of all enzymes in the region used for antibody production and derived from MPK9. The MPK9 phosphopeptide synthesized is shown above the alignment. The N-terminal K was added to allow efficient coupling to carrier protein KLH. (B) Dot–blots of peptides spotted to the membrane after coupling to BSA. Membranes were probed with crude serum diluted 5000-fold with the addition of each peptide as labelled.

**Figure S6. Predicted Arabidopsis thaliana MPK9 structure.**

The AlphaFold structure for MPK9 corresponding to UniProt ID Q9LV37 was downloaded from the AlphaFold Protein Structure Database (https://alphafold.ebi.ac.uk/). PyMol was used to visualize the structure and residues after tyrosine 376 were hidden for image clarity. Marked in pink is the activation loop, defined by DFG and APE, showing how the putative PP1 docking RVxF SLIM (light blue) is surface exposed. The activation loop TDY motif threonine (T) and tyrosine (Y) are presented with side chains shown (dark blue).

**Figure S7. Percentage germination after 7 days in the presence of ABA.**

ABA sensitivity of Nössen*, rlph2-1*, *rlph2-2*, Col-0 and 35S::RLPH2-Ctap line (RLPH2 OE). Seeds were stratified 4d at 4°C on MS agar medium without ABA (control) or in the presence of 1, 5 and 20 µM ABA. The germination rate was scored after 7d on light at 21°C. Results represent germination means ± SE of three replicates of fifty seeds. Asterisks indicate statistical differences between Nössen and *rlph2* mutants or between Col-0 and the RLPH2 OE line, as determined by One-way ANOVA with post hoc Tukey’s test (P < 0.05).

**Figure S8. Percentage germination in the presence of PAC after 7d.**

PAC sensitivity of Nössen, *rlph2-1*, *rlph2-2*, Col-0 and 35S::RLPH2-Ctap line (RLPH2 OE). Seeds were stratified 4d at 4°C on MS agar medium without PAC (control) or in the presence of 1, 3, and 10 µM PAC. The germination rate was scored after 7d on light at 21°C. Results represent germination means ± SE of three replicates of fifty seeds. Asterisks indicate statistical differences between Nössen and *rlph2* mutants or between Col-0 and RLPH2 OE line, as determined by One-way ANOVA with post hoc Tukey’s test (P < 0.05).

**Figure S9. Conserved region of C-terminal tails of Arabidopsis D-group MPKs.**

The D-group MPKs were aligned and the conserved region among the C-terminal tails extracted and presented. The most conserved residues are highlighted in red.

**Table S1. Plant proteins with pTxpY motifs extracted from Plant PTM Viewer (www.psb.ugent.be/webtools/ptm-viewer/).**

**Table S2: All Tyr phosphorylated peptides identified by Mascot and Scaffold Q+S.**

## References and Notes

1. D. Kerk, C. White-Gloria, J. J. Johnson, G. B. Moorhead, Eukaryotic-like Phosphoprotein Phosphatase (PPP) enzyme evolution: interactions with environmental toxins and regulatory proteins. Biosci Rep 43, (2023).

2. D. L. Brautigan, S. Shenolikar, Protein Serine/Threonine Phosphatases: Keys to Unlocking Regulators and Substrates. Annu Rev Biochem 87, 921–964 (2018).

3. N. Sugiyama et al., Large-scale phosphorylation mapping reveals the extent of tyrosine phosphorylation in Arabidopsis. Mol Syst Biol 4, 193 (2008).

4. R. G. Uhrig, A. M. Labandera, G. B. Moorhead, Arabidopsis PPP family of serine/threonine protein phosphatases: many targets but few engines. Trends Plant Sci 18, 505–513 (2013).

5. D. Kerk et al., The origin and radiation of the phosphoprotein phosphatase (PPP) enzymes of Eukaryotes. Sci Rep 11, 13681 (2021).

6. D. Kerk, G. Templeton, G. B. Moorhead, Evolutionary radiation pattern of novel protein phosphatases revealed by analysis of protein data from the completely sequenced genomes of humans, green algae, and higher plants. Plant Physiol 146, 351–367 (2008).

7. A. M. Labandera, R. G. Uhrig, K. Colville, G. B. Moorhead, K. K. S. Ng, Structural basis for the preference of the Arabidopsis thaliana phosphatase RLPH2 for tyrosine-phosphorylated substrates. Sci Signal 11, (2018).

8. R. G. Uhrig, A. M. Labandera, J. Muhammad, M. Samuel, G. B. Moorhead, Rhizobiales-like Phosphatase 2 from Arabidopsis thaliana Is a Novel Phospho-tyrosine-specific Phospho-protein Phosphatase (PPP) Family Protein Phosphatase. J Biol Chem 291, 5926–5934 (2016).

9. M. A. Schumacher, J. L. Todd, A. E. Rice, K. G. Tanner, J. M. Denu, Structural basis for the recognition of a bisphosphorylated MAP kinase peptide by human VHR protein Phosphatase. Biochemistry 41, 3009–3017 (2002).

10. W. Zhang et al., The MPK8-TCP14 pathway promotes seed germination in Arabidopsis. Plant J 100, 677–692 (2019).

11. M. Group, Mitogen-activated protein kinase cascades in plants: a new nomenclature. Trends Plant Sci 7, 301–308 (2002).

12. Y. Y. Zhang, Z. Q. Mei, J. W. Wu, Z. X. Wang, Enzymatic activity and substrate specificity of mitogen-activated protein kinase p38alpha in different phosphorylation states. J Biol Chem 283, 26591–26601 (2008).

13. T. Tanoue, T. Yamamoto, E. Nishida, Modular structure of a docking surface on MAPK phosphatases. J Biol Chem 277, 22942–22949 (2002).

14. A. J. Bardwell, E. Frankson, L. Bardwell, Selectivity of docking sites in MAPK kinases. J Biol Chem 284, 13165–13173 (2009).

15. W. Peti, R. Page, Molecular basis of MAP kinase regulation. Protein Sci 22, 1698–1710 (2013).

16. S. K. Nagy et al., Activation of AtMPK9 through autophosphorylation that makes it independent of the canonical MAPK cascades. Biochem J 467, 167–175 (2015).

17. G. R. Lampard, D. L. Wengier, D. C. Bergmann, Manipulation of mitogen-activated protein kinase kinase signaling in the Arabidopsis stomatal lineage reveals motifs that contribute to protein localization and signaling specificity. Plant Cell 26, 3358–3371 (2014).

18. S. K. Nagy et al., Activation of AtMPK9 through autophosphorylation that makes it independent of the canonical MAPK cascades. Biochemical Journal 467, 167–175 (2015).

19. M. Bollen, W. Peti, M. J. Ragusa, M. Beullens, The extended PP1 toolkit: designed to create specificity. Trends Biochem Sci 35, 450–458 (2010).

20. M. S. Choy et al., Understanding the antagonism of retinoblastoma protein dephosphorylation by PNUTS provides insights into the PP1 regulatory code. Proc Natl Acad Sci U S A 111, 4097–4102 (2014).

21. J. Mergner et al., Mass-spectrometry-based draft of the Arabidopsis proteome. Nature 579, 409–414 (2020).

22. G. W. Templeton et al., Identification and characterization of AtI-2, an Arabidopsis homologue of an ancient protein phosphatase 1 (PP1) regulatory subunit. Biochem J 435, 73–83 (2011).

23. S. P. Lyons et al., A Quantitative Chemical Proteomic Strategy for Profiling Phosphoprotein Phosphatases from Yeast to Humans. Mol Cell Proteomics 17, 2448–2461 (2018).

24. S. S. Taylor, A. P. Kornev, Protein kinases: evolution of dynamic regulatory proteins. Trends Biochem Sci 36, 65–77 (2011).

25. T. K. Mohanta, P. K. Arora, N. Mohanta, P. Parida, H. Bae, Identification of new members of the MAPK gene family in plants shows diverse conserved domains and novel activation loop variants. BMC Genomics 16, 58 (2015).

26. Z. Y. Zhang et al., Substrate specificity of the protein tyrosine phosphatases. Proc Natl Acad Sci U S A 90, 4446–4450 (1993).

27. D. R. Alessi et al., Inactivation of p42 MAP kinase by protein phosphatase 2A and a protein tyrosine phosphatase, but not CL100, in various cell lines. Curr Biol 5, 283–295 (1995).

28. I. Nasa, S. F. Rusin, A. N. Kettenbach, G. B. Moorhead, Aurora B opposes PP1 function in mitosis through phosphorylation of the conserved RVxF binding motif in PP1 regulatory proteins Sci Signal, (2018).

29. R. G. Uhrig, P. Schlapfer, B. Roschitzki, M. Hirsch-Hoffmann, W. Gruissem, Diurnal changes in concerted plant protein phosphorylation and acetylation in Arabidopsis organs and seedlings. Plant J 99, 176–194 (2019).

30. R. G. Uhrig et al., Diurnal dynamics of the Arabidopsis rosette proteome and phosphoproteome. Plant Cell Environ 44, 821–841 (2021).

31. J. Leymarie et al., Role of reactive oxygen species in the regulation of Arabidopsis seed dormancy. Plant Cell Physiol 53, 96–106 (2012).

32. P. Willems et al., The Plant PTM Viewer 2.0: in-depth exploration of plant protein modification landscapes. J Exp Bot, (2024).

