## Supplemental files for "Phospho-proteomics identifies D-group MAP kinases as substrates of the Arabidopsis tyrosine phosphatase RLPH2"

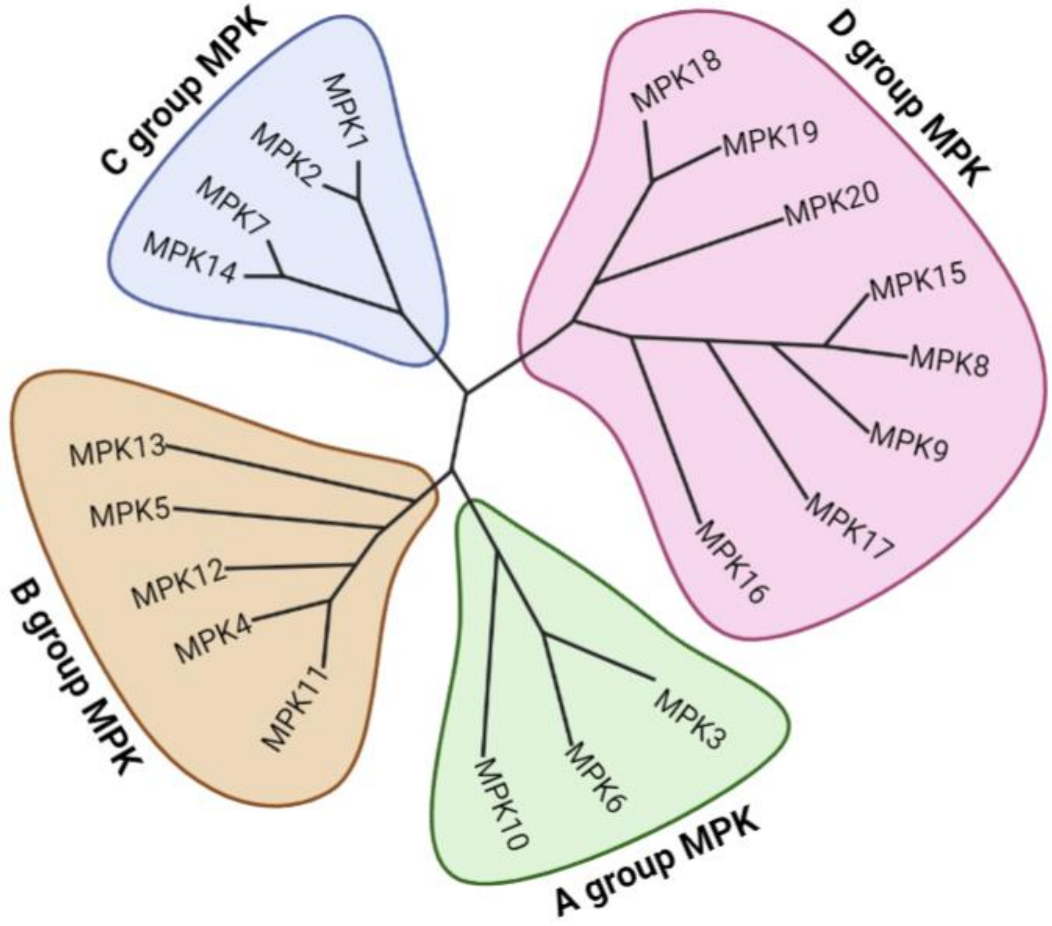


**Supplemental Figure S1. Phylogenetic analysis of the 20 MPK proteins in Arabidopsis thaliana.** Groups A, B, C and D are labeled with the corresponding MPKs present in each group. Sequence Viewer 8.0 was used to align the sequences and generate the phylogenetic tree.

| Group | MPK | Sequence |
|---|---|---|
| A | MPK3 | DFGLARPTS----ENDFMTEYVVTRWYRAPE |
|  | MPK6 | DFGLARVTS----ESDFMTEYVVTRWYRAPE |
|  | MPK10 | DFGLARATP----ESNLMTEYVVTRWYRAPE |
| B | MPK4 | DFGLARTKS----ETDFMTEYVVTRWYRAPE |
|  | MPK5 | DFGLARTTS----ETEYMTEYVVTRWYRAPE |
|  | MPK11 | DFGLARTKS----ETDFMTEYVVTRWYRAPE |
|  | MPK12 | DFGLARTTS----DTDFMTEYVVTRWYRAPE |
|  | MPK13 | DFGLARTSN----ETEIMTEYVVTRWYRAPE |
| C | MPK1 | DFGLARASNT---KGQFMTEYVVTRWYRAPE |
| | MPK2 | DFGLARTSNT---KGQFMTEYVVTRWYRAPE |
| | MPK7 | DFGLARTSQG---NEQFMTEYVVTRWYRAPE |
|  | MPK14 | DFGLART------YEQFMTEYVVTRWYRAPE |
| D | MPK8 | DFGLARVSFNDAPTAIFWTDYVATRWYRAPE |
| | MPK9 | DFGLARVSFNDAPSAIFWTDYVATRWYRAPE |
| | MPK15 | DFGLARVSFNDAPTAIFWTDYVATRWYRAPE |
| | MPK16 | DFGLARVAFNDTPTAIFWTDYVATRWYRAPE |
| | MPK17 | DFGLARVSFTDSPSAVFWTDYVATRWYRAPE |
| | MPK18 | DFGLARVAFNDTPTTVFWTDYVATRWYRAPE |
| | MPK19 | DFGLARVSFNDTPTTVFWTDYVATRWYRAPE |
| | MPK20 | DFGLARVAFNDTPTTIFWTDYVATRWYRAPE |

**Supplemental Figure S2: Alignment of Arabidopsis thaliana MPK activation loops.** Protein kinase activation loops are defined by the sequences DFG and APE. In MPKs activation is achieved by phosphorylation of a TXY motif (green). Arabidopsis MPKs form 4 phylogenetically distinct groups as indicated (A-D) with only the D-group have an aspartate (D) between the T and Y of the TXY motif, while groups A-C have a E in this position (*11*). The D-group enzymes also possess an insert in the loop that conforms to a protein phosphatase one (PP1) binding RVXF SLiM (blue).

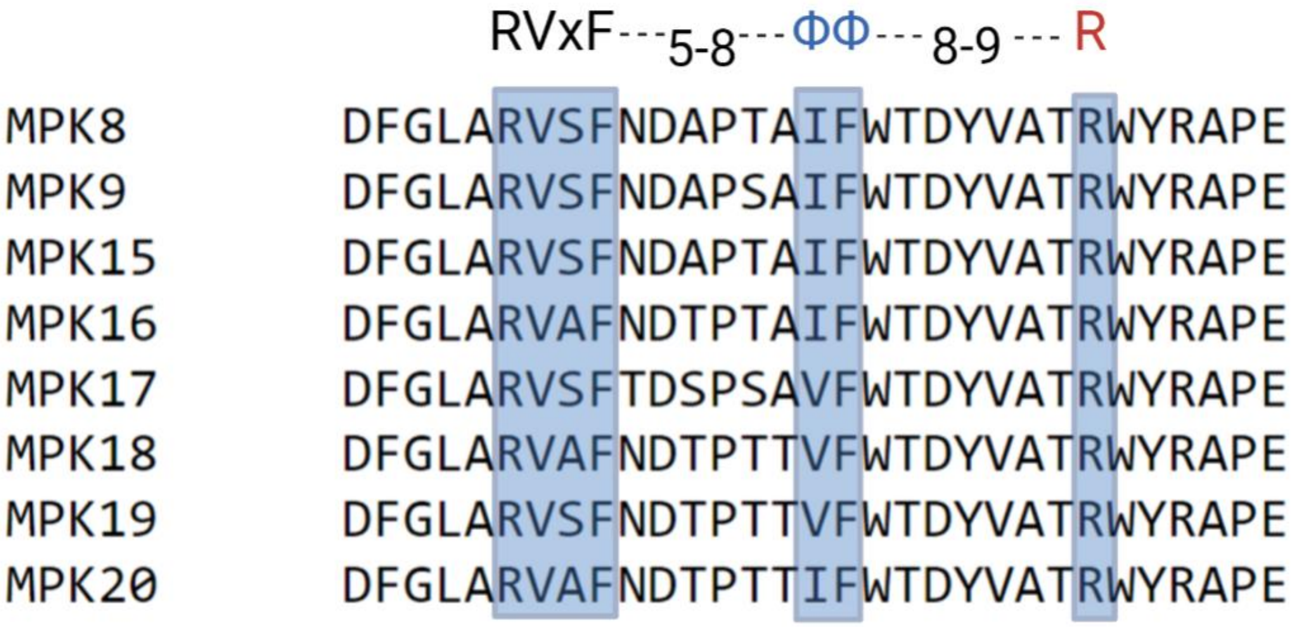


**Supplemental Figure S3: D-group MPKs have an extended RVXF motif**. The activation loops of the D group MPKs as defined by DFG and APE. Highlighted in blue are the amino acids associated with the extended RVXF motif, which starts with RVXF, then 5-8 amino acids C-terminal two hydrophobic residues (Val, Ile or Phe), and then 8-9 amino acids C-terminal an arginine (R). Each D-group enzyme conforms to this sequence motif further supporting the idea that these protein kinases recruit PP1.

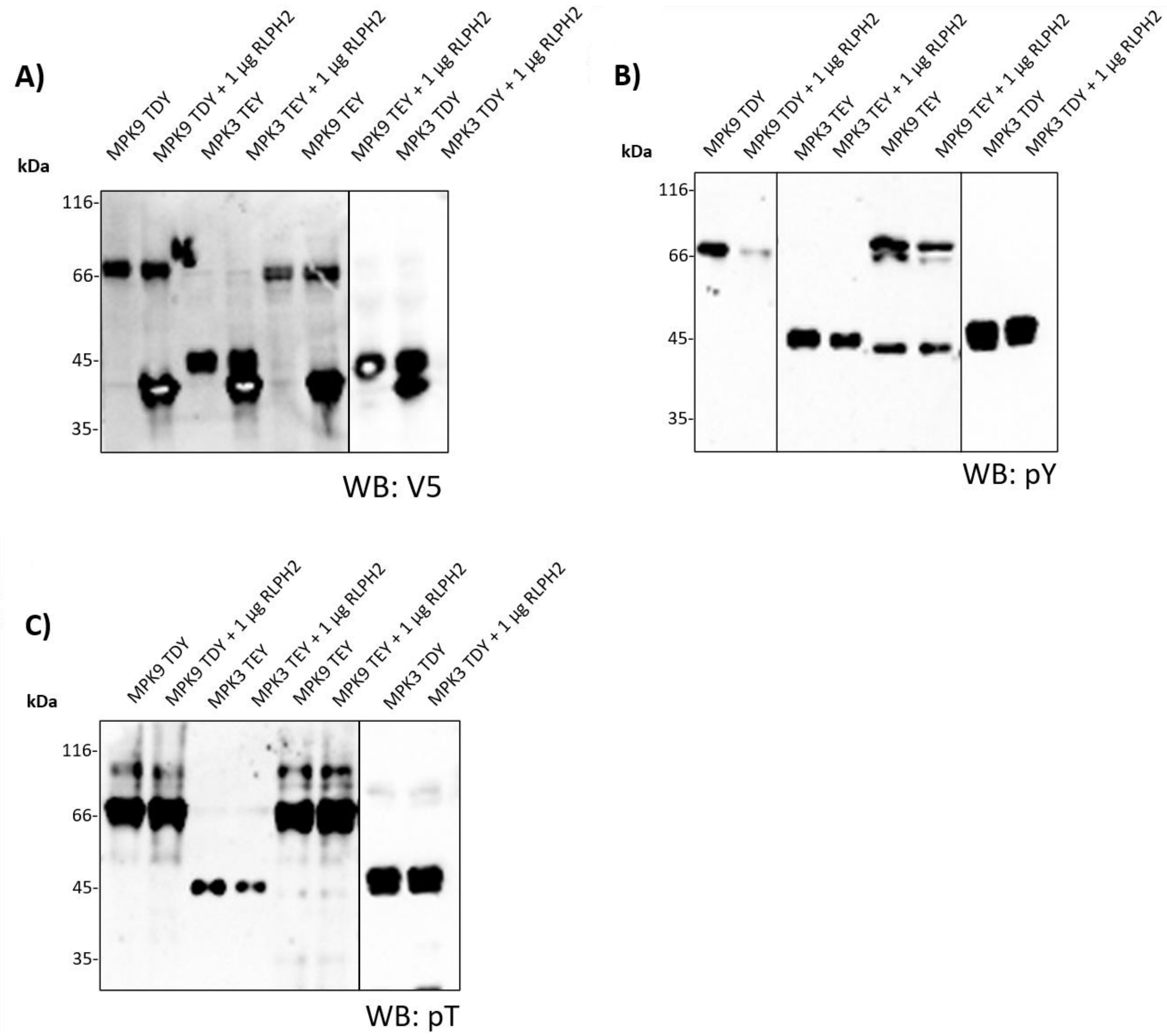


**Supplemental Figure S4: RLPH2 solely dephosphorylates pY of MPK9 TDY, but not MPK9 TEY, MPK3 TEY or MPK3 TDY**. MPKs were dually phosphorylated in their activation loops (see methods) and treated with or without purified RLPH2. Western blotting was performed to assess the MPK phospho-status with either anti-pY or pT antibodies. (**A**) Anti-V5 immunoblot demonstrating equal loading. (**B** & **C**) Anti-phosphotyrosine and anti-phosphothreonine immunoblots, respectively.

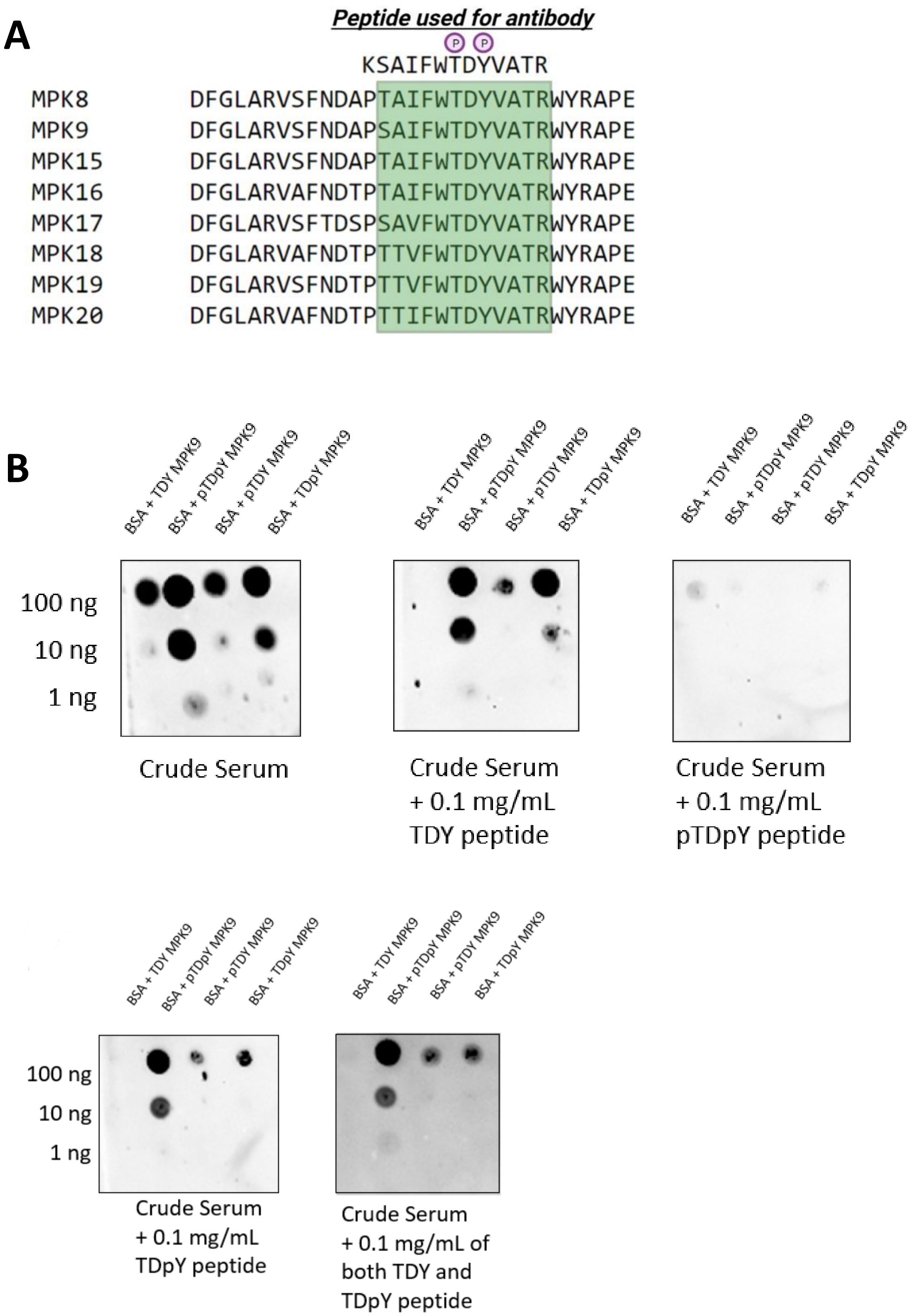


**Supplemental Figure S5**: **Characterization of MPK9 TDY phospho-specific antibodies.** (**A**) Alignment of D-group MPK activation loops highlighting the sequence relationship (boxed green) of all enzymes in the region used for antibody production and derived from MPK9. The MPK9 phospho-peptide synthesized is shown above the alignment. The N-terminal K was added to allow efficient coupling to carrier protein KLH. (**B**) Dot–blots of peptides spotted to the membrane after coupling to BSA. Membranes were probed with crude serum diluted 5000-fold with the additions of each peptide as labelled.

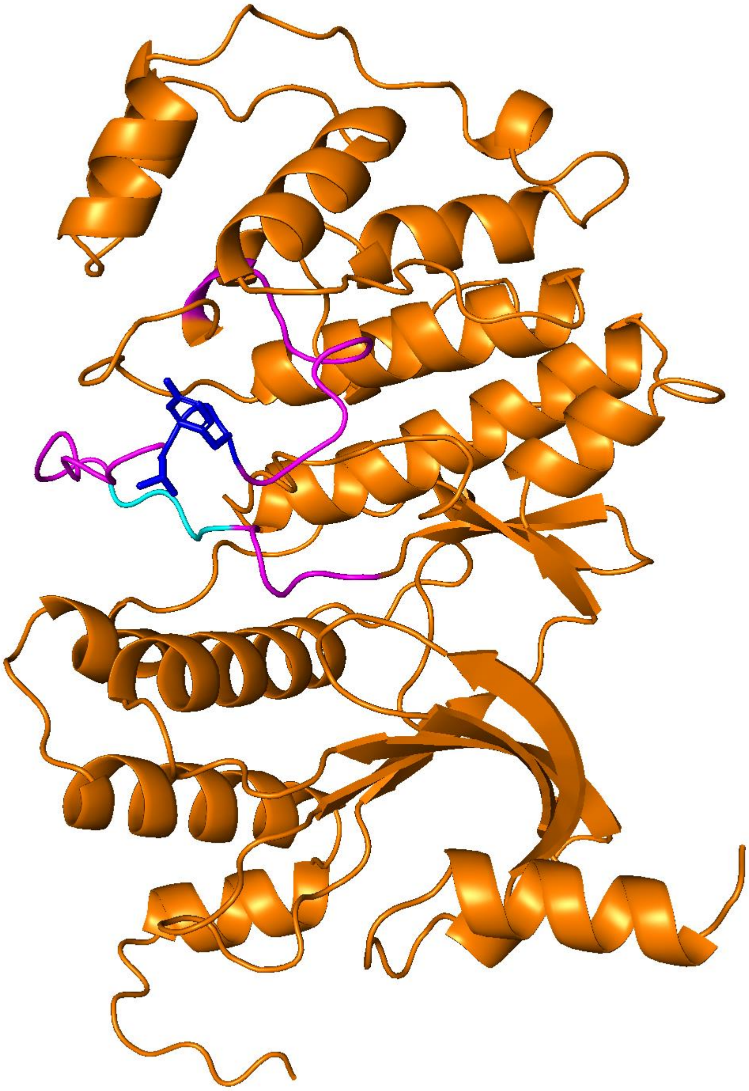

**Supplemental Figure S6: Arabidopsis thaliana MPK9 structure**. The AlphaFold structure for MPK9 corresponding to UniProt ID Q9LV37 was downloaded from the AlphaFold Protein Structure Database (https://alphafold.ebi.ac.uk/). PyMol was used to visualize the structure and residues after tyrosine 376 were hidden for image clarity. Marked in pink is the activation loop, defined by DFG and APE, showing how the putative PP1 docking RVxF SLIM (light blue) is surface exposed. The activation loop TDY motif threonine (T) and tyrosine (Y) are presented with side chains shown (dark blue).

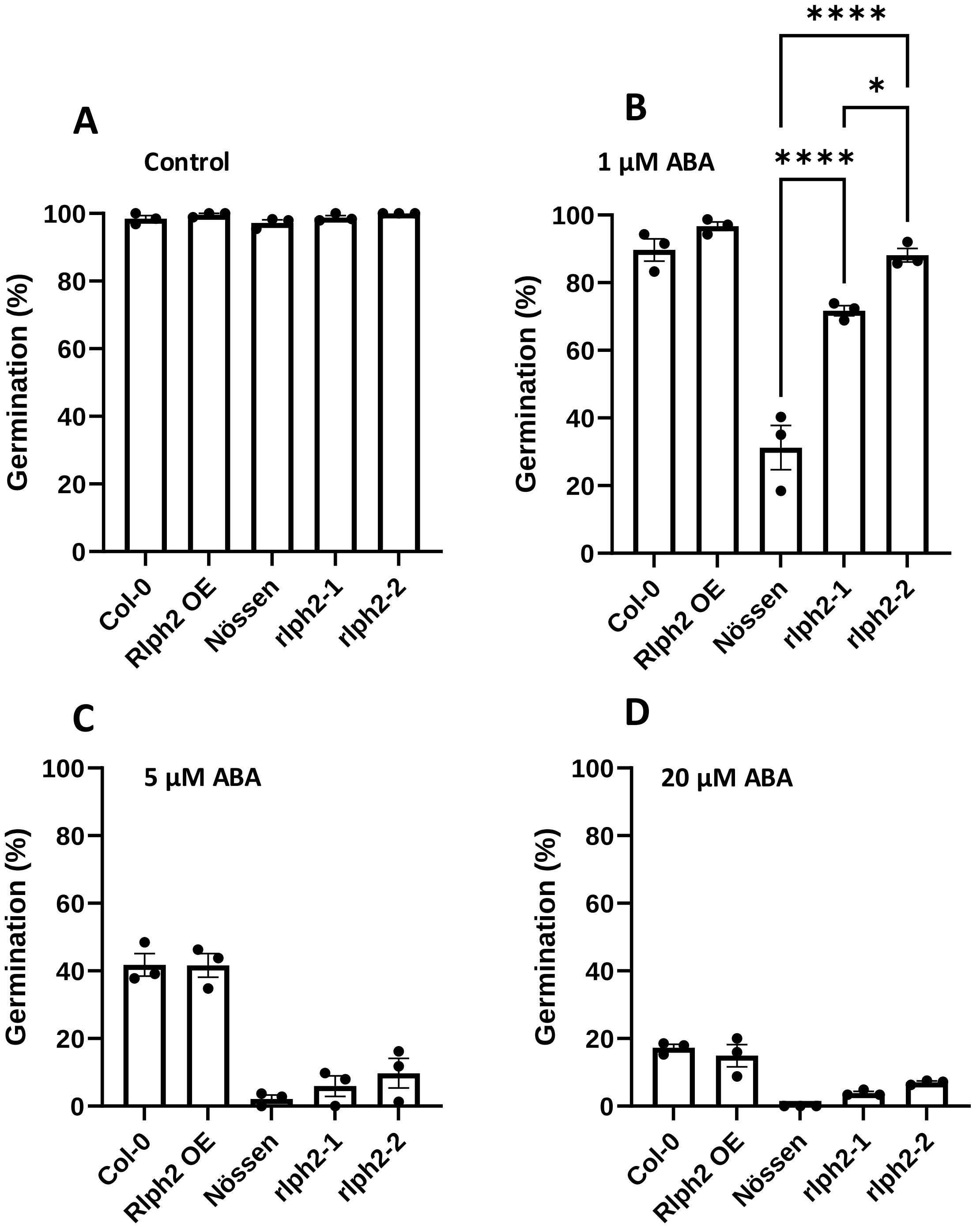


**Supplemental Figure S7. Percentage germination after 7 days in the presence of ABA.** ABA sensitivity of Nössen, *rlph2-1*, *rlph2-2*, Col-0 and 35S::RLPH2-Ctap line (RLPH2 OE). Seeds were stratified 4d at 4°C on MS agar medium without ABA (control) or in the presence of 1, 5 and 20 µM ABA. The germination rate was scored after 7d on light at 21°C. Results represent germination means ± SE of three replicates of fifty seeds. Asterisks indicate statistical differences between Nössen and *rlph2* mutants or between Col-0 and the RLPH2 OE line, as determined by One-way ANOVA with post hoc Tukey's test ($P < 0.05$).

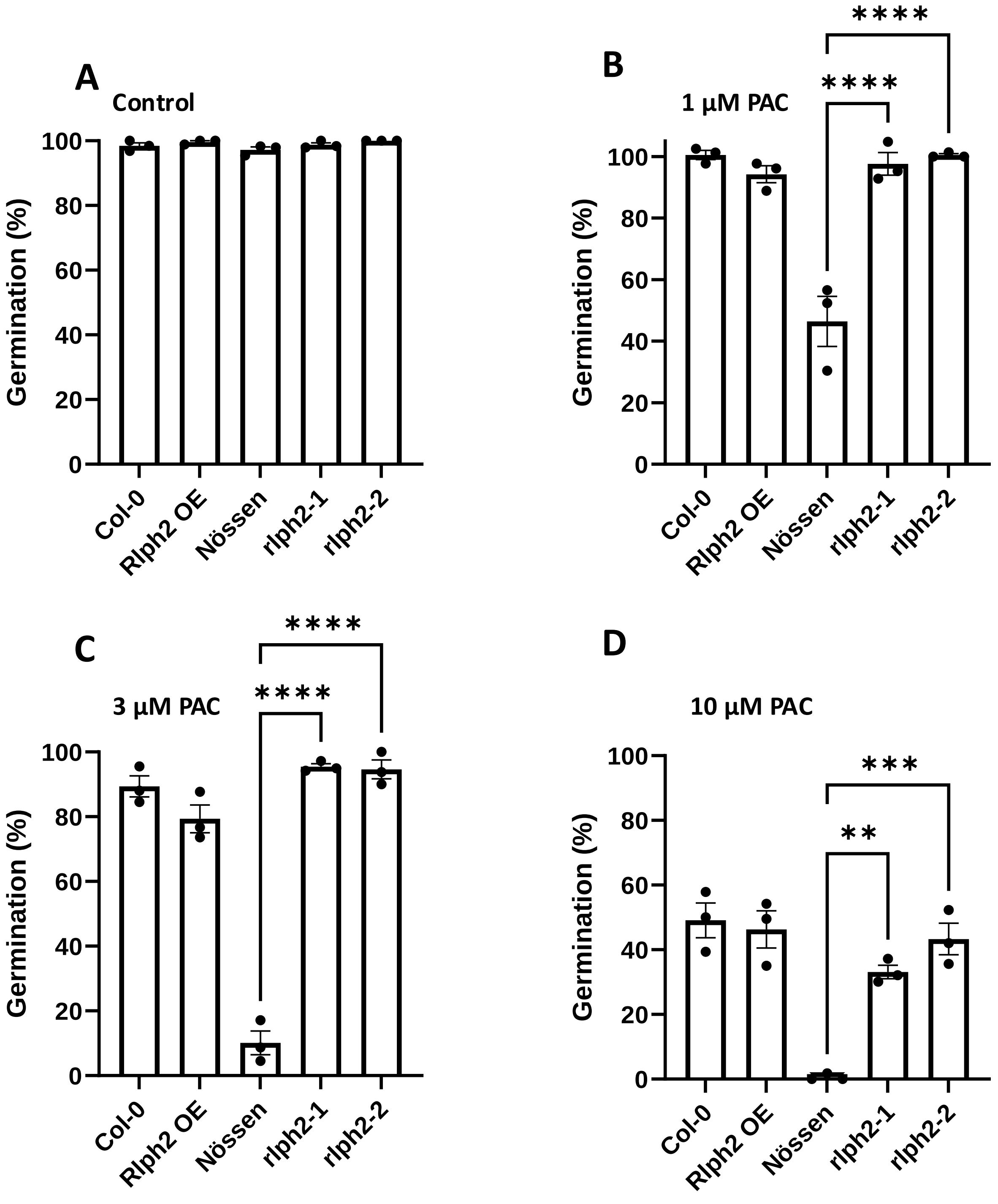


**Supplemental Figure S8. Percentage germination in the presence of PAC after 7d.** PAC sensitivity of Nössen, *rlph2-1*, *rlph2-2*, Col-0 and 35S::RLPH2-Ctap line (RLPH2 OE). Seeds were stratified 4d at 4°C on MS agar medium without PAC (control) or in the presence of 1, 3, and 10 µM PAC. The germination rate was scored after 7d on light at 21°C. Results represent germination means ± SE of three replicates of fifty seeds. Asterisks indicate statistical differences between Nössen and *rlph2* mutants or between Col-0 and RLPH2 OE line, as determined by One-way ANOVA with post hoc Tukey's test ($P < 0.05$).

```
MPK8    DQLS--FMYPSGVDRFKRQFAHLEENQGKPGAAGGGRSTALHRHHASLPR
MPK15   NQLS--FMYPSGVDRFRRQFAHLEENQGP-----GGRSNALQRQHASLPR
MPK9    EQTS--FMYPSGVDRFKRQFAHLEENYGK-----GEKGSPLQRQHASLPR
MPK17   ENINSHFLYPSGVDQFKQEFARLEEHNDDEE---EHNSPPHQRKYTSLPR
MPK16   EPTN--FMYPSAVEHFKKQFAYLEEHYKNG----TSHNPPERQQHASLPR
MPK18   EGSN--FVYPSAIGHLRQQFTYLEENSSRN-----GPVIPLERKHASLPR
MPK19   EGSS--FLYPSAIGHLRKQFAYLEENSGKS-----GPVIPPDRKHASLPR
MPK20   DKAS--FLYPSAVDQFRRQFAHLEENSGKT-----GPVAPLERKHASLPR
```

**Supplemental Figure S9: Conserved region of C-terminal tails of *Arabidopsis thaliana* D-group MPKs.** The D-group MPKs were aligned and the conserved region among the C-terminal tails extracted and presented. The most conserved residues are highlighted in red.
